# Single Cell Spatial Analysis and Biomarker Discovery in Hodgkin Lymphoma

**DOI:** 10.1101/2023.05.24.542195

**Authors:** Alexander M Xu, Aixiang Jiang, Tomohiro Aoki, Alicia Gamboa, Lauren Chong, Anthony Colombo, Yifan Yin, Joseph Lownik, Katsuyoshi Takata, Monirath Hav, Christian Steidl, Akil Merchant

## Abstract

The biology of tumors is suffused with spatial interactions, such as tumor-immune signaling through localized cytokine/ligand secretion, cell-cell contacts, and checkpoint ligand/receptor signaling. Hodgkin Lymphoma (HL) can serve as a study paradigm for tumor microenvironment (TME) architecture as the defining pathological feature is the scarcity of the malignant Hodgkin and Reed Sternberg (HRS) cells, leaving a diverse and predominantly immune cell rich tumor microenvironment (TME) with complex tumor-immune interactions. Previous studies have identified TME features that are prognostic and predictive, however these studies did not consider the entirety of TME cellular ecosystems, including precisely defined immune cell subsets with opposing inflammatory and immune-suppressive effects, as a determinant for differential clinical course of HL patients. Here we use Imaging Mass Cytometry (IMC) with 42 antibody markers to profile tumors from 93 patients with HL. Our cohort consists of relapsed/refractory HL with matched diagnostic and relapsed biopsies, and we present a bioinformatic pipeline to profile 10 major cell lineages and their subtypes including spatial interaction mapping. Our pipeline identifies putative biomarker candidates with a focus on “rosettes” – local aggregates of immune cells around single tumor cells. In addition to validating existing biomarkers centered on CD68+ macrophages, GranzymeB+CD8+ T cells, and others in HL, we propose new biomarkers based on localized interactions between HRS cells and aggregating CD4+ and CD8+ T cells and macrophages involving the immune checkpoints PD1/PDL1, LAG3, and Galectin9. This study serves as a broad tissue imaging resource for multi-timepoint biopsies in HL, and a computational resource and pipeline for users of IMC and other multiplexed imaging studies to perform tissue analysis and biomarker candidate testing with any tissue type.

## BACKGROUND

Spatial analytics is an essential tool in the clinical oncology repertoire and a common diagnostic method for diagnosing cancer and predicting outcomes when used in pathology and immunohistochemical (IHC) staining ^1–3^. Image interpretation is performed by trained pathologists, who integrate many factors including cell and nuclear morphology, staining intensity, and spatial context of cells^4–6^. These observations guide clinical decisions, based on the tumor structure, risk factors, and treatment options available^7, 8^. By comparing a single tissue stain to the historical record of tissue accrued over time, patterns in the tissue can be associated with patient survival and clinical outcomes after treatment. These patterns, along with clinical and molecular data with similar predictive power, are called biomarkers.

Tumors exhibit a wide range of host immune response which is reflected in the numbers of immune cells which are found surrounding and infiltrating the malignant cells. Hodgkin Lymphoma (HL), an extreme example in a spectrum of diseases that feature an immune-rich tumor-microenvironment (TME), is composed of a multitude of non-malignant cell types educated by malignant Hodgkin and Reed-Sternberg (HRS) cells, which are relatively isolated and comprise <10% of the tissue^9–11^. The pattern and structure of the TME has been used to classify Hodgkin lymphoma into histological subtypes – nodular sclerosis, mixed cellularity, lymphocyte-rich and lymphocyte depleted; however, these spatial classifications have little prognostic impact and are not relevant to initial treatment selection. Instead, prognoses are made using clinical variables such as International Prognostic Score^12, 13^ or positron-emission tomography^14–16^. Standard treatment of primary HL consists of doxorubicin, bleomycin, vinblastine, and dacarbazine (ABVD)^17^, and recent clinical trials confirmed the benefit of an additional CD30 targeting treatment, brentuximab vedotin, in advanced HL^18–20^. Despite these treatment advancements, about 20-30% of patients still experience relapse within a year or are refractory to treatment^21, 22^. About half of relapsed/refractory HL patients can achieve long term remission after high dose chemotherapy followed by autologous stem cell transplantation.

Previous work in HL has that suggested specific cell types such as FoxP3+ T cells^23^, macrophages^24, 25^, and many others^26^ have prognostic value, but the spatial context of these cells was overlooked. Recently, HL-specific spatial patterns have been explored in new ways using multiplexed imaging technologies. PDL1 overexpression is common in HL tumor cells due to chromosomal translocations, and the recent use of multiplexed imaging studies has shown that PDL1+ macrophages and monocytes are found closer to PDL1+ HRS cells, where they interact with PD1+CD4+ T cells^27, 28^. Other checkpoints such as CTLA4 follow their own spatial patterns of expression in the TME^29^. Roemer et al. reported a study of 108 cases where a spatial biomarker, based on the relative MHCI expression on HRS cells to their immediate neighbors, predicted the results of chemotherapy^30^. The observation that HRS cells recruit other immune cells, especially CD4+ T cells, to form aggregates called “rosettes”, suggests that spatial analysis of HRS-immune cell neighbor interactions is particularly significant^31, 32^.

Here we present one of the largest collections of HL reported and analyzed with highly multiplexed, single cell, spatial protein analysis with subcellular resolution. A total of 93 HL patients were analyzed, including 73 patients with relapsed/refractory disease and an additional 20 patients with no relapse^33^. For each patient that relapsed, paired diagnostic pretreatment and relapse tumor tissues were profiled. Our formalin-fixed paraffin embedded (FFPE) tissue data was analyzed using Imaging Mass Cytometry (IMC) to measure 35 proteins at 1 μm resolution^34–36^. We present a detailed analytical pipeline to process and analyze these image data. We performed deep phenotyping and spatial analysis to describe the tumor architecture and functional protein expression profiles of major cell types, and we emphasized hyper-local protein expression by extracting patterns of immune checkpoint ligand-receptor co-expression between HRS and immune cells. Finally, we present a computational pipeline to propose biomarker candidates using IMC data, which have both protein and spatial components. We explored specific biomarkers using Cox survival analysis, and we also provide a tool to generate biomarker combinations based on a variant of the least absolute shrinkage and selection operator (LASSO), again tailored to different spatial tumor contexts.

## RESULTS

### Hodgkin Lymphoma Preliminary Analysis

We studied a cohort of 93 total HL patients (Table 1) comprising 261 regions of interest (ROIs) assembled on tumor microarrays (TMAs) with up to 2 timepoints per patient and up to 2 ROIs per timepoint when available (average 1.76). We selected a cohort containing samples from patients with variable relapse status, including 20 non-relapsed patients and 73 relapsed patients, of which 26 relapsed within 1 year (‘early relapse’) and 47 relapsed later than 1 year (‘late relapse’) (FIG 1A). All relapsed patients had matched samples from diagnostic pretreatment and relapse time points, enabling comparisons over time. The panel used for HL analysis consisted of primary phenotyping lineage markers, secondary lineage markers, and functional or inducible markers (FIG 1B). Tumor cells as defined by CD30^37^ along with immune cell types were first identified, and cell subtypes were then selected based on checkpoint expression. Proteins reported as biomarkers or potential therapeutic targets are denoted in Figure 1B^23–26, 30, 33, 38–56^. Our cohort was designed to identify relapse-specific biological features and develop biomarker assays to improve clinical decision making. Our specific marker panel reflected the demonstrated and growing importance of the TME and immune checkpoints in HL treatment.

**Figure 1.**
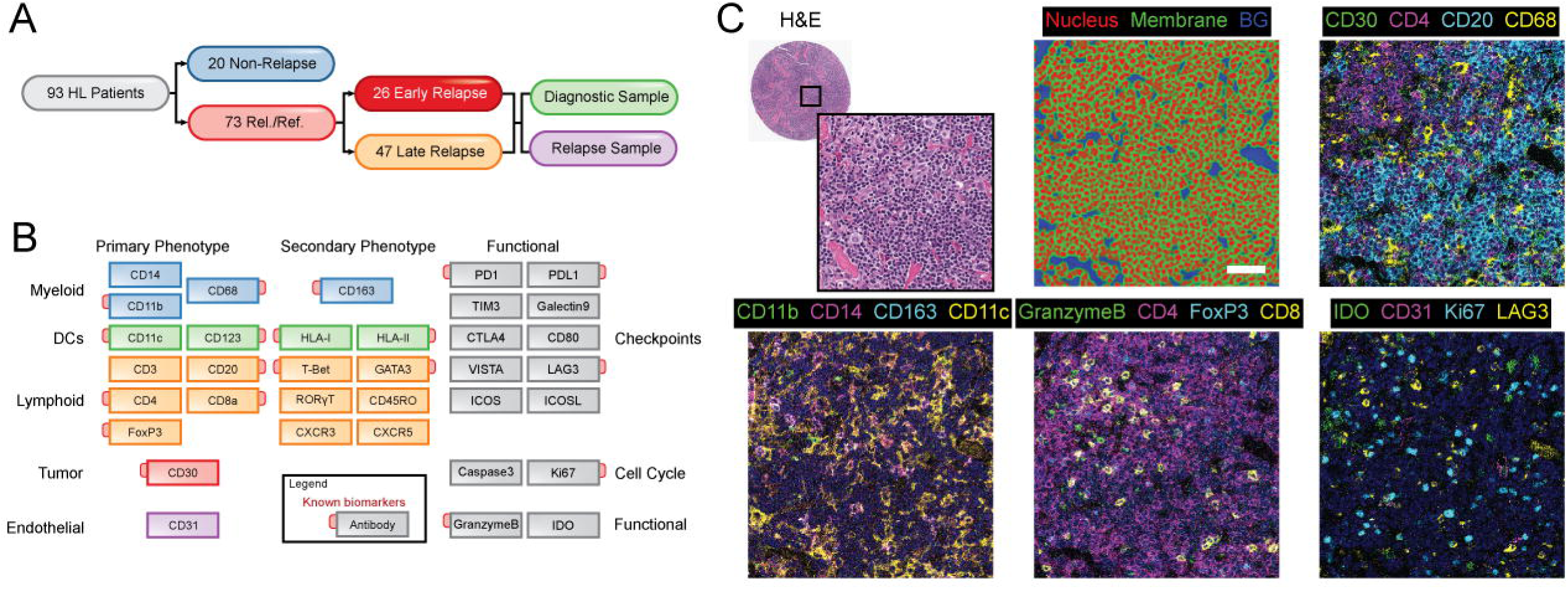
IMC experimental design. A. Consort diagram showing relapse status (non-relapse, late relapse, early relapse) and timepoint (diagnostic and relapse). B. A panel of 35 antibodies was used to image tissue. Antibodies were organized into three tiers to identify major cell phenotypes, their subtypes, and functional subsets of cell phenotypes. C. Sample images for an ROI are shown, showing the H&E and inset image (top left), segmentation results using ilastik (top middle), tumor and general immune phenotypes (top right), myeloid and dendritic cell markers (lower left), lymphocyte markers (lower middle), and markers indicating immune cell function (lower right). Scale bar – 25 μm.

**Table 1.** Patient characteristics of cohort.

**Table 2.** Description of IMC antibody panel.

We analyzed 7.05M cells total using our spatial analytic pipeline (SFIG1A), for an average of 27,002±9,754 cells per sample, and 46,984±22,085 cells per patient timepoint. Average marker expression in cells was approximately normally distributed across ROIs (SFIG1B). Representative images showed a heterogenous immune microenvironment with classic HL morphology, consisting of large HRS cells embedded in a matrix of immune cells in the TME (FIG 1C). Segmentation partitioned each image into nuclei, cytoplasm/membrane, and background areas used to identify single cells, and protein expression patterns specific to lymphocytes, myeloid cells, and other functional markers are shown. All subsequent analysis was performed on single cells using average protein expression of each marker.

### HL Immune Phenotyping

We identified 10 major cell types using hybrid hierarchical and manual metaclustering (SFIG2A-E): T cells (CD4+, CD8+, Treg), B cells, macrophages and other myeloid lineage cells, conventional and plasmacytoid dendritic cells, endothelial cells, and HRS tumor cells (FIG 2A). In a UMAP projection (downsampled to 10%), cell clusters were crowded due to the densely-packed lymphoma tissue and imperfect segmentation (FIG 2B). Across patients, major cell types were heterogeneously distributed (SFIG2F,G). The largest proportion of cells were T cells (∼31%) split into CD4+ (16%), CD8+ (11%), and Treg subtypes (6%), B cells (23%), and myeloid cells (20%, 6.5% macrophages). Dendritic cells (7% conventional, 2.5% plasmacytoid), tumor cells (7.5%), and endothelial cells (4.5%) comprised the remainder of the tissue (FIG 2C). The 7.5% abundance of HRS tumor cells we observed was likely due to our emphasis on relapsed/refractory tumors, which are enriched in HRS cells^57, 58^. Out of 7.05M cells, ∼6% were found in clusters that were not well-defined by a single phenotype. Of these, 382k expressed multiple canonical cell phenotyping markers, which were denoted as “mixed” cells (SFIG2D,E). For these difficult-to-define cells, each cell was labeled with as many descriptors as applicable (i.e. CD4+ T cell, macrophage), which were not mutually exclusive to account for the possibility that two cells were overlapping in the same section of the tissue slice.

**Figure 2.**
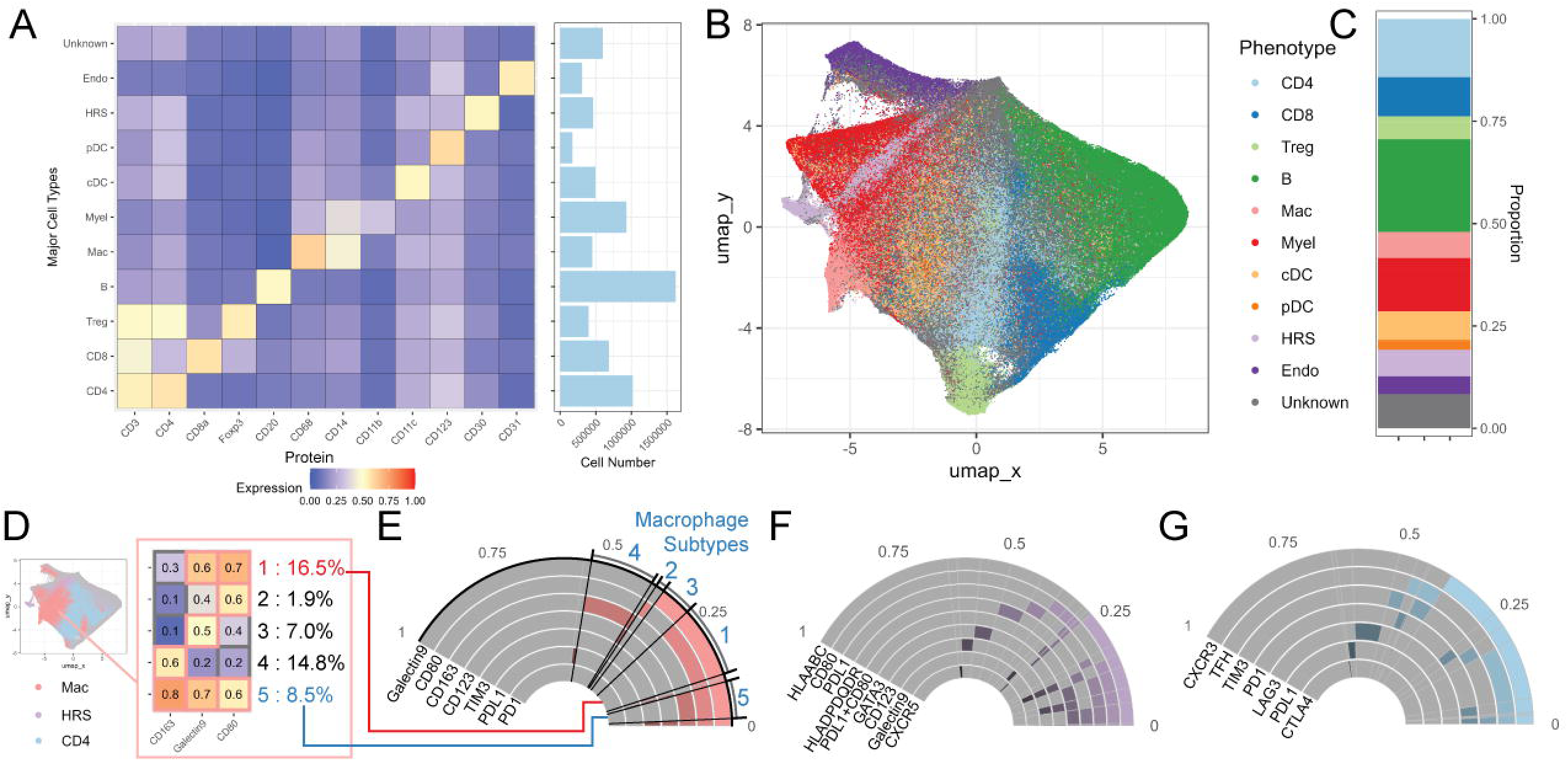
HL phenotyping. A. A total of 10 major cell types were identified, with their mean expression heatmap and absolute proportions displayed. Cells that were unable to be classified within any of the categories were labeled “Unknown”. B. A UMAP was generated of all cells with major cell types colored. Unknown cells were often found between clusters. C. Total relative proportions of all cell types are shown. D. Sample strategy for clustering within major cell types is shown using 5 macrophage subtypes. The subtypes express combinations of CD163, Galectin9, and CD80, labeled 1-5. E. The coexpression patterns and relative proportions of macrophage subtypes are shown with subtypes 1-5 from panel D highlighted. F. Coexpression pattern of HRS cell subtypes. G. Coexpression pattern of CD4+ T cell subtypes.

Within immune cell types, additional clustering was performed to identify relevant subtypes (SFIG3A,B). Similar to the mixed cell process, we adopted a strategy of labeling cells with as many subtypes as applicable, where a subtype was defined by 1-2 protein markers (FIG 2D-G, SFIG3C). To illustrate this strategy, a heat map of a macrophage subset expressing CD163, Galectin9, CD80, and their combinations is shown (FIG 2D). The proportion of these 5 clusters relative to all macrophages is shown in Figure 2E, where macrophages co-expressing Galectin9 and CD80 (label 1) or all three markers (label 5) were common while CD80+ only macrophages (label 2) were relatively rare. Ki67+ proliferative cells comprised 8% of all cells but 22% of HRS and 16% of Tregs (SFIG3D-E). HRS cells (11%), macrophages (16%), and CD8+ T cells (14%) were more apoptotic (Caspase3+) compared to all cells (8%, SFIG3D,F). Among T cell subtypes, CD45RO expression, which denotes a memory subtype, was increased in PD1+ and TIM3+ T cells, and decreased in LAG3+ T cells (SFIG3G). Our phenotyping approach was single-marker-focused to facilitate downstream biomarker analysis, where subtypes such as PDL1+ HRS cells can be recalled easily but complex phenotypes were still identified via clustering.

### Spatial Analysis

To quantify the organization of the heterogenous HL TME, we calculated spatial metrics from a single-cell perspective. Using a modified nearest neighbor (NN) analysis^59^ (SFIG4A), we calculated spatial enrichment scores to describe every cell’s spatial enrichment with the 10 major cell types. Homotypic interactions – cell interactions with other cells of the same type – were most common, especially among B cells (FIG 3A). Beyond homotypic interactions, tumor cells interact more with CD4+ T, cDC, and myeloid cells, and less with B, CD8+ T, and endothelial cells, consistent with existing literature^9, 60^. A correlation plot summarizes the significance of the spatial enrichments observed (SFIG4B). The spatial enrichment scores were used for two purposes: 1) as input variables for biomarker candidates; and 2) to organize single cells into distinct local microenvironments (similar to “niches” or “neighborhoods”)^61, 62^.

**Figure 3.**
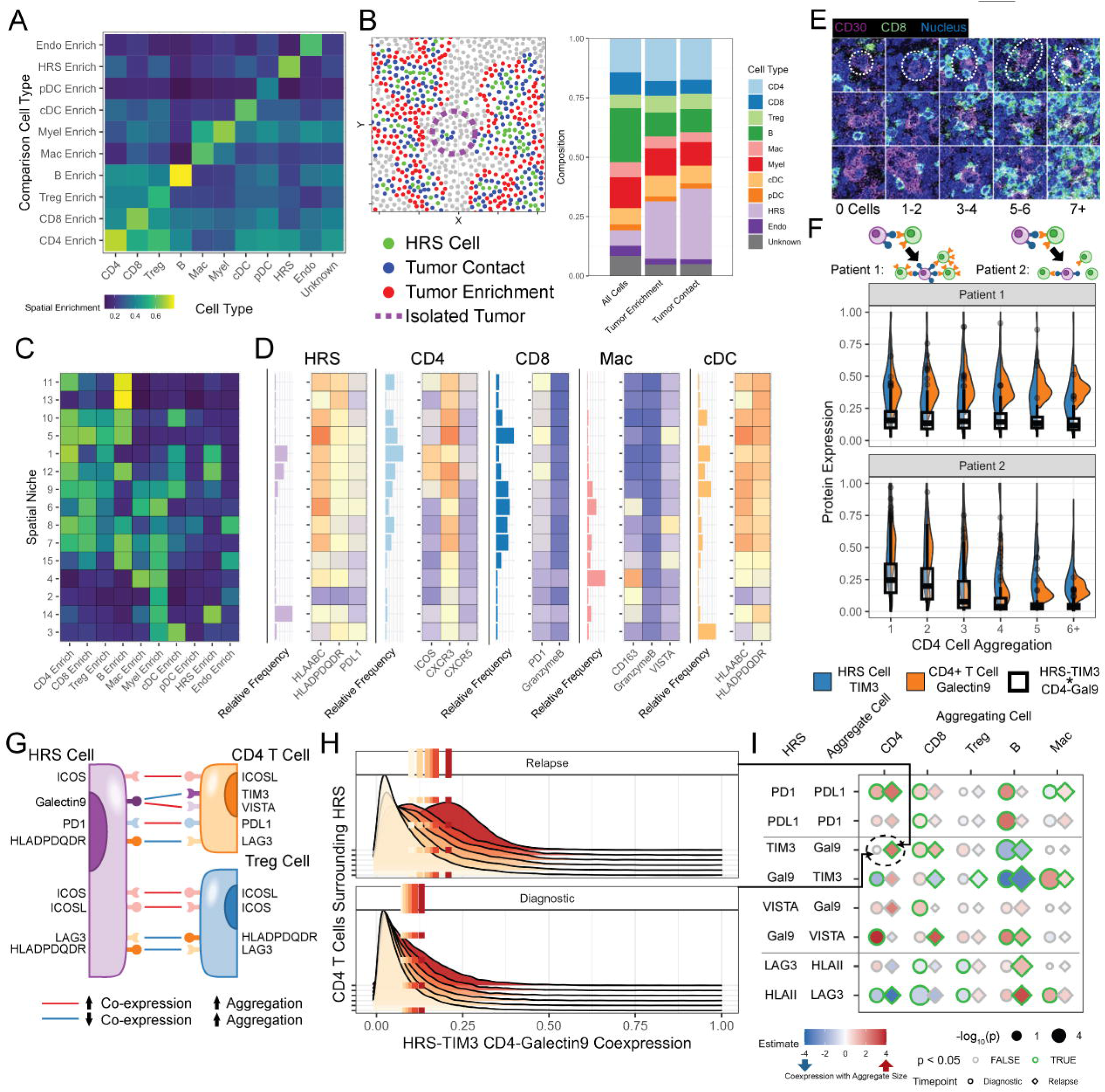
Spatial architecture of HL. A. The average spatial enrichment of each cell type for every other cell type. Spatial enrichments were not reciprocal due to differences in their abundance, i.e. rare pDCs were spatially enriched with CD4+ T cells but not vice versa. B. Sample image of “digital biopsies” using thresholds of 0.5 and 10 μm for HRS spatial enrichment and nearest neighbor (NN) distance measurements, and cell proportions of tumor-enriched and tumor-contact regions. C. Using k-means clustering of spatial enrichment measurements with k=15, spatial cell niches defined. D. The frequency and cell-type specific expression of select markers in HRS, CD4+, CD8+ T cells, macrophages and cDCs in each of the 15 spatial niches. E. Aggregation of specific immune cells around HRS cells was seen at different extents, from 0 aggregating cells to 7+ aggregating cells. The aggregate size was defined as the number of cells with < 15μm centroid-to-centroid distance relative to a central cell. F. The ligand receptor co-expression of HRS cells and aggregating immune cells is shown as a function of the number of aggregating cells, and the strength and direction of the correlation varies. In patient 1, HRS TIM3, average CD4+ T cell Galectin9 expression, and their co-expression product are independent of the number of aggregating CD4+ T cells. In patient 2, the ligand-receptor pair co-expression is inversely correlated with aggregate size. G. Ligand receptor pair co-expression was measured and used to model rosette size for CD4+ and CD8+ T cells, with significant (p<0.05) measurements highlighted and colored by the direction of the association. H. The average coexpression of HRS-TIM3/CD4-Galectin9 is strongly associated with aggregate size in relapsed tumor samples, but not diagnostic samples. I. The significance of ligand-receptor coexpression measurements associated with aggregate size in diagnostic/relapse samples shows a distinct pattern for each cell type.

We used the spatial enrichment scores to isolate comparable regions of analyzed tissue between patients. Due to tissue heterogeneity and the relatively small size of each patient’s analyzed tissue (∼1 mm^2^), a frequent concern in IMC analysis of TMAs is the generalizability of comparisons across patients in the face of heterogeneity^63–66^. To mitigate this, we isolated cells in similar HRS cell-enriched microenvironments to perform a “digital biopsy” by selecting cells with a tumor spatial enrichment score > 0.5 (tumor region) or a nearest-neighbor distance to HRS cells of < 10μm (tumor contact, FIG 3B). As expected, both types of tumor-enriched regions had increased HRS and CD4+ T cell proportions and reduced CD8+ T cell and B cell proportions^31, 32^. The primary difference between the two biopsies was their treatment of isolated tumor cells, where were ignored by the tumor region biopsy (purple dashed circle).

We also used the spatial enrichment scores to identify different types of local TME/niches using k-means clustering^67^. Each cluster represents local regions of cells with enriched or depleted spatial interactions between specified other cell types. There was no well-defined elbow to determine the optimal cluster number^68^, so a cluster number of 15 was used based on existing literature (SFIG4C)^61, 62^. The unique spatial architecture of HL is reflected in the composition of spatial clusters (FIG 3C). HRS tumor cells were found in clusters enriched in CD4+/Treg cells (Cluster 1), T and B cells (Cluster 12), or myeloid cells (Cluster 14). Organized cell structures such as residual follicles (B cell rich clusters 13 and 11) were found along with B and T cell mixed zones (Clusters 1, 5, 7, 8, 9, 10, 11, 12). No tumor-only clusters were observed due to the low density of HRS cells. The composition of the niches led to niche-specific protein expression on major cell types (FIG 3D). In lymphocyte-rich clusters, increased expression levels of HLA proteins were observed in HRS and dendritic cells (Clusters 6, 7, 9, 10, 12). Among macrophages, CD163 expression was negatively correlated with dendritic cell proximity (Clusters 1, 9, 10, 12 vs 2, 4, 6, 14, 15). Among clusters with significant HRS tumor presence (>5%), some HRS immune checkpoint proteins were correlated with proximity to B cells (CTLA4) and proximity to myeloid cells (PD1). We next focused on a specific spatial pattern example that was more localized and 5characteristic to HL.

Hyper-local CD4+ T cell clustering around HRS cells has long been observed in HL and has been historically defined as a “rosette”^31, 32, 69^. We hypothesized that cell aggregation around HRS cells, whether as CD4+ T cells or a different immune cell type, was the result of the local balance of inflammatory and suppressive signals, and furthermore, the extent of aggregation and functional state of aggregated cells may be linked to clinical outcomes. We first quantified cell aggregation by counting the number of cells of selected subtypes (CD4+, CD8+, Treg, Macrophage, B, FIG 3E, SFIG4D) in immediate proximity (<15 μm) of HRS cells. We found that aggregates formed with higher frequency than random chance in T cells and macrophages using a random replacement model (SFIG4E-F)^70^. Since aggregation was non-random, we sought to determine if it was dependent on functional cellular interactions.

We hypothesized that correlating the expression of receptor-ligand pairs on HRS and aggregating cells would help us understand how rosettes were formed and maintained^71–76^. We measured the average cell-type specific protein expression on the aggregating cells and the central HRS cell as a function of the number of aggregating cells (FIG 3F). By modeling the contributions of all positive receptor-ligand signaling pairs expression levels to aggregate size, we identified protein pairs with both positive (FIG 3G, HRS-ICOSL/Treg-ICOS)^77^ and negative (HRS-HLADPDQDR/CD4-LAG3)^78^ correlations to aggregate size in diagnostic samples. Here, a significant correlation indicated that co-expression increased/decreased with increased aggregate size. These patterns provide evidence for a mechanism of positive/negative feedback in local microenvironments based on spatial cell arrangement.

To examine if these protein expression and aggregation patterns had clinical significance we compared pretreatment diagnostic and posttreatment relapsed samples. Certain ligand-receptor pairs, such as HRS-ICOSL/Treg-ICOS and HRS-HLADPDQDR/CD8-LAG3 (SFIG4G) were found across both diagnostic and relapse samples, and could represent universal aspects of HL biology independent of chemotherapy treatment. We observe significant reorganization in local PD1/PDL1, HLADPQDR/LAG3, and Galectin9 signaling between diagnostic and relapse HL tumor biology (FIG 3H-I). Galectin9 was differentially associated with aggregation between diagnostic and relapse samples in many instances, including on CD4+ T cells (FIG 3H). PD1/PDL1 signaling increased with CD8+ T cell and B cell aggregates at diagnosis but not relapse. We observe significant PD1/PDL1 signaling with macrophage aggregation at relapse but not diagnosis in this hyper-local setting, for a similar result as other studies concluded at longer length scales^27^. HLADPDQDR/LAG3 was negatively associated with CD8+ and Treg aggregates in diagnostic samples only, before emerging as a significant association in B cell aggregates upon relapse. These ligand-receptor results, along with HRS protein-only or aggregating cell-only results (SFIG4H-I) show the evolution of local environments of HRS tumor cells between diagnosis and relapse and the pathways potentially shaping these environments.

### Biomarker Validation Using IMC

Although most patients with HL are cured with initial chemotherapy, for the subset of patients who are refractory or experience early relapse the overall prognosis is much poorer as they are less likely to be cured with salvage therapies like autologous bone marrow transplant^79^. To better identify biomarkers that could identify these high-risk patients, we focused our analysis on diagnostic samples associated with early, late, or no relapse. We used IMC analysis of our cohort to identify and validate protein and cell patterns linked to survival and patient outcomes. We called these measured cell percentages or expression levels “biomarker candidates” and determined their significance using overall survival as the clinical endpoint initially. First, biomarkers from literature were compared to equivalent IMC biomarker candidates. Biomarkers using sample-level proportions of cells such as CD68+ macrophages^24^ were readily validated with IMC (FIG 4A). More granular biomarkers based on cell-specific expression, such as GranzymeB+CD8+ T cells^39, 40^ (FIG 4B) and PDL1+ HRS cells^49^ (FIG 4C), were also validated. Immunohistochemistry-based biomarkers often use complex scoring strategies. Roemer et al proposed one such scoring strategy based on MHCI/II expression on the tumor cell relative to its neighbor cells^30^. We used aggregation analysis to automate the identification and stratification of MHC-I-negative cells and validated this biomarker (FIG 4D, SFIG5A).

**Figure 4.**
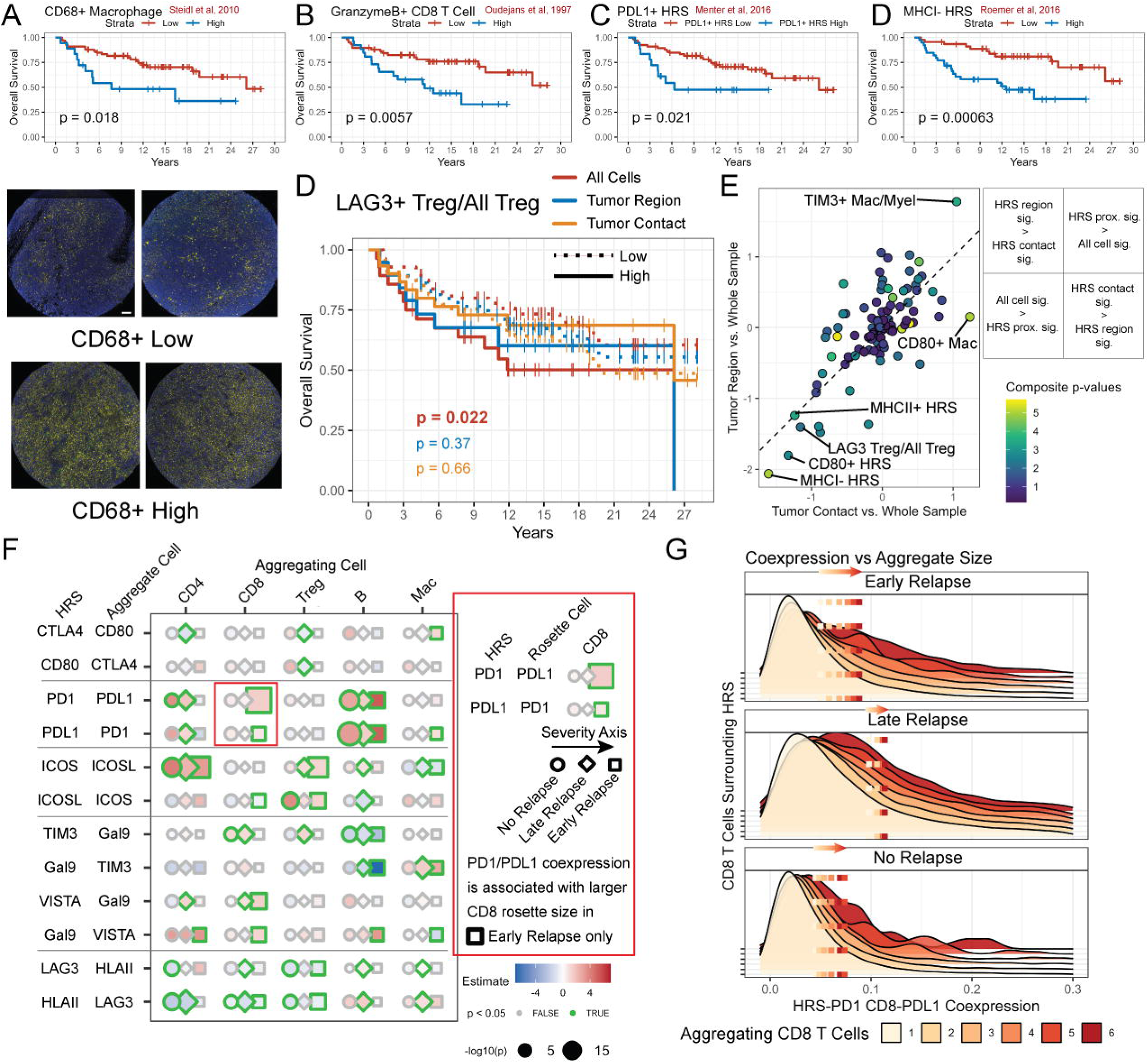
Biomarker Testing and Validation. A. Kaplan-Meier (KM) survival curve for CD68+ cell proportion and 4 sample images of CD68+ low and high samples. Scale bar – 100 μm. B-D. KM curves for GranzymeB+CD8+ T cells, PDL1+ HRS cells, and MHC-HRS cells. E. The fraction of LAG3+ Tregs out of all Tregs was used as a biomarker in all cells or one of two digital biopsies (tumor region – HRS spatial enrichment >0.5, tumor contact – HRS nearest neighbor distance < 10 μm). KM curves for the digital biopsies are compared, with a significance survival difference observed in all cells but neither biopsy. E. The relative significance of biomarker candidates in tumor region or tumor contact digital biopsies relative to the whole image is shown in diagnostic samples predicting overall survival. Biomarkers fall in one of 4 quadrants, with HRS-based biomarkers found in the bottom left (more significant in whole images), some immune biomarkers in the top right (more significant in tumor-enriched biopsies). Color indicates the product of p-values across the three spatial categories (all cells, tumor region, tumor contact, –log10). F. Receptor-ligand coexpression patterns associated with aggregate size in diagnostic samples by relapse status. G. HRS-PD1/CD8-PDL1 coexpression vs. CD8+ T cell aggregate size by relapse status, with a statistically significant association in early relapse patients only.

We hypothesized that spatial organization may be predictive of HL outcomes, and that it can be used to add additional context to biomarkers. Significant changes in cell composition were observed between patients based on their relapse status (no relapse, late relapse, early relapse) and sample timepoint (diagnostic, relapse). CD8+ T cell and B cell aggregates increased in late relapse patients (SFIG5B, log scale, Tukey’s HSD test), especially in their relapse samples. Among spatial niches, a myeloid cluster was significantly more abundant in non relapse patients while HRS-enriched spatial niches were enriched in relapse patients (SFIG5C-D).

We used digital biopsies to compare tumor-enriched regions and tumor-contacting cells between patients to refine biomarkers of overall survival in diagnostic samples to be less sensitive to intrapatient heterogeneity. LAG3+ Tregs as a percentage of total Tregs^69, 78^ were one such biomarker that was predictive only when all cells were used but not in either digital biopsy, thus suggesting that subtype of Tregs distant from the tumor were the predictive elements (FIG 4D). On the other hand, CD80+ Macrophages^29, 80^ were one of many biomarkers that became significant only in tumor-enriched regions (FIG 4E).

Macrophage^55, 81^ and HRS^30, 82^ cell subtypes were the most frequently significant biomarkers along with established biomarkers such as GranzymeB+ CD8+ T cells. This remained true when additional clinical outcomes such as disease-specific survival, 1^st^ failure-free survival, 2^nd^ failure-free survival, and postBMT failure-free survival were considered (SFIG5E). Biomarker analysis for categorical clinical variables (EBV, MHCI, MHCII, Early Relapse, No Relapse) showed that CD8+ T cells and HRS subtypes were the most significant biomarker candidates, and HRS candidates were more spatially dependent (SFIG5F). We also evaluated single cell variants for significant biomarker candidates (p<0.05 after Benjamini-Hochberg). For such subtypes, i.e. PD-L1+CD8+ T cells, we tested the equivalent cell-type-specific protein expression (PD-L1 expression on single CD8+ T cells, SFIG5G). Most candidates were significant at both sample and single cell levels except for those tracking negative expression (HLAABC-HRS, CXCR5-HRS) or subtype percentages (GranzymeB+CD8+ T cell vs total CD8+ T cell).

We found ligand-receptor protein expression in local HRS immune aggregates to be dependent on the relapse status of patients (FIG 4F, SFIG5H-L). For example, while HRS-PD1/CD8-PDL1 signaling was significantly associated with CD8+ T cell aggregate size across all diagnostic samples (FIG 3I), we found this to be significant in early relapse diagnostic samples only after stratifying diagnostic samples by patient relapse status (FIG 4G). In contrast, PD1/PDL1 in CD4+ T cell aggregates was significant in non-relapse or late relapse samples only. Other early relapse patterns included HRS-ICOSL/Mac-ICOS, HRS-Galectin9 and TIM3 on B cells and Macrophages, and HRS-Galectin9 and VISTA on CD4+, CD8+, B cells, and macrophages. Significant patterns in non relapse patients included HRS-TIM3/CD8-Galectin9, and LAG3/HLADPDQDR signaling on CD4+ T cells. Finally, some patterns were selectively significant in late-relapse patients only, including CTLA4/CD80, ICOS/ICOSL expression on B cells, and LAG3/HLADPDQDR on B cells and macrophages. These patterns may reflect the ability of immune cells to regulate HRS behavior and feedback mechanisms of signaling that result in increased immune recruitment. Furthermore, these patterns show that a ligand-receptor signaling pathway may have different consequences for tumor relapse status depending on the immune cell type involved.

### Hodgkin Lymphoma Biomarker Discovery

The biomarkers explored above were focused on cell-type-specific expression, which can involve as many as 3-5 proteins in the context of a particular cell type. We searched for novel combinations of sample-level cell proportions that predicted overall survival using a dimensional reduction and variable selection strategy (LASSO_plus) based on the least absolute shrinkage and selection operator (LASSO), single variable selection, and stepwise variable selection. In diagnostic samples, LASSO_plus selected HLADPDQDR HRS cells with higher expression than their neighbors and PDL1+CD80+ Macrophages as significant biomarker candidates, with GranzymeB+CD8+ T cells, Galectin9+ Macrophages and Myeloid cells, and other CD80+ cells as marginal candidates (FIG 5A). The LASSO_plus strategy enables a comprehensive search for broader potential biomarkers without the need for manual intervention and facilitates the construction of a model based on selected variables. This is achieved through the utilization of our automated R function, LASSO_plus, which is accessible in our new R package csmpv (https://github.com/ajiangsfu/csmpv).

**Figure 5.**
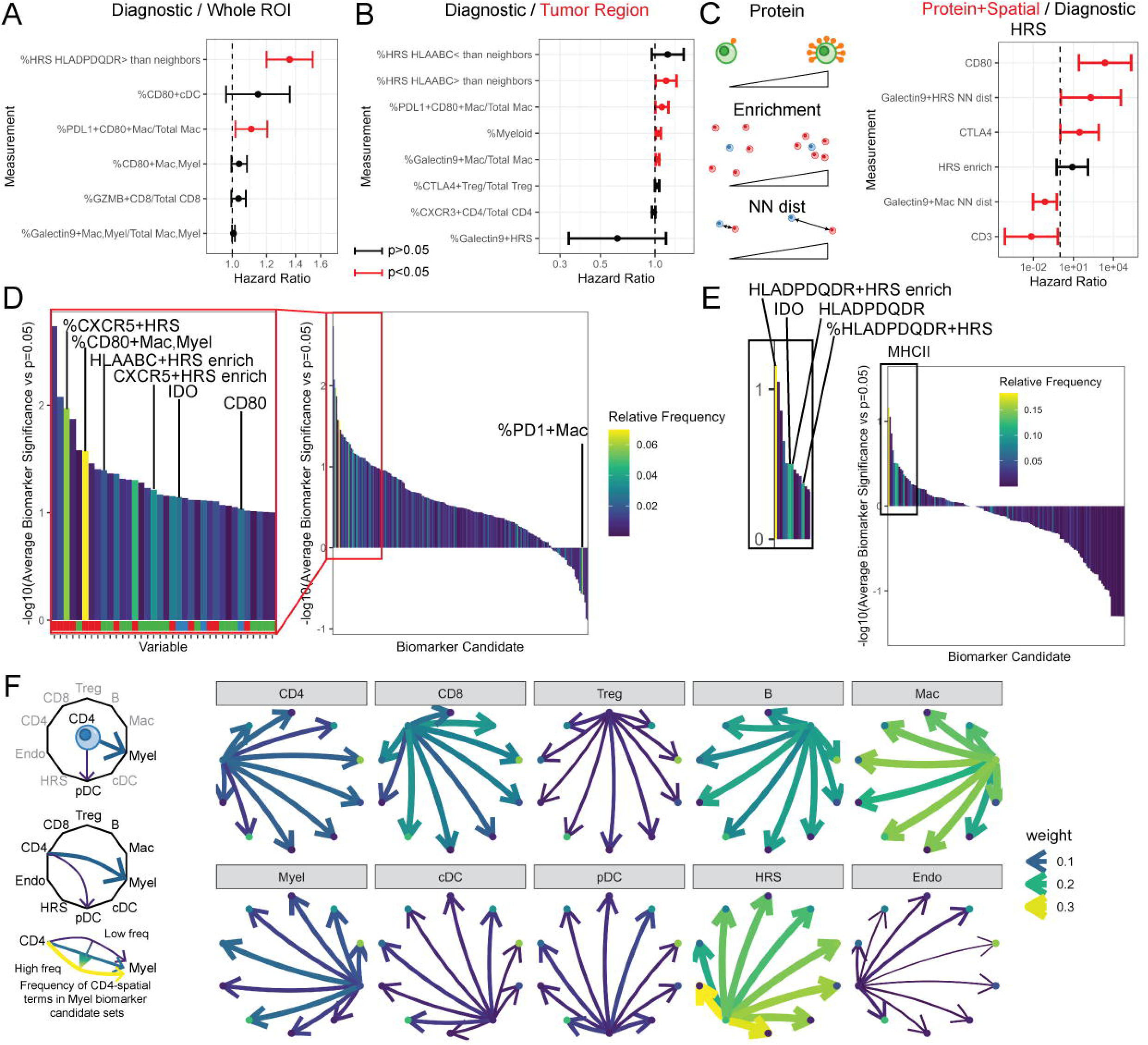
Biomarker Discovery with LASSO_plus. A. Biomarker candidate set of cell proportions identified using diagnostic samples with LASSO_plus. B. Biomarker candidate set identified using tumor region digital biopsy. C. Combined protein and spatial biomarker candidate set. D. Waterfall plot showing the type of biomarker candidate vs. the average significance of the candidate relative to p=0.05 for all major cell types, digital biopsies, timepoints, and survival conditions. Frequently appearing cell type proportions, proteins, and spatial measurements are highlighted. E. Waterfall plot for biomarker candidates generated to predict MHCII status, with common HLADPDQDR-related candidates as well as proteins such as IDO. F. Among spatial biomarker candidates, the frequency of LASSO_plus selection of spatial biomarkers is plotted as a function of cell type, showing the predictive power of specific cell-type spatial relationships.

Our spatial and clinical data enabled additional biomarker candidate testing using LASSO_plus. Using the digital biopsy to isolate tumor regions revealed a different set of biomarker candidates in diagnostic samples, including HLAABC-HRS cells and Myeloid cells (FIG 5B). Relapse samples reflect an altered tumor microenvironment, which yielded different biomarker candidates, including Galectin9+ HRS and pDCs, PDL1+ CD4+ T cells, and HLADPDQDR+ HRS cells (SFIG6A-B). Cell specific average protein expression and spatial enrichments and distances were explored as a cell-centric biomarker format using LASSO_plus instead of an image-level proportion biomarker in diagnostic and relapse samples (FIG 5C, SFIG6C). These two biomarker formats are related (see Galectin9+ macrophage significance in both cases), but the cell centric biomarker may be less sensitive to changes in cell proportions caused by heterogeneous sampling. This pipeline can also be used to explore biomarkers measured on other cell types such as macrophages (SFIG6D).

We searched for proteins, spatial metrics, and cell proportions with the highest average significance (geometric mean vs. p=0.05), and highest relative frequency among all LASSO_plus biomarker candidate sets and plotted them as waterfall-style plots (FIG 5D, SFIG6E-G). Among proteins, IDO, Galectin9, CD3, and CD80 were an order of magnitude more significant than p=0.05 on average (geometric mean of p<0.005), while IDO, CD80, and CD3 (along with CXCR5, CXCR3, and CD20) appeared in more than 350 sets (>2% of all possible). Spatial enrichment of Tregs, HLAABC+ HRS, CD163+ macrophages, PDL1+CD4+ and HRS cells, and Galectin9+ Macrophages were the most significant biomarker candidates while HRS, B cells, CXCR5+ HRS and B cells, HLAABC+ HRS cells, and macrophages were the most frequent.

A similar biomarker candidate discovery analysis for clinical factors (EBV status, MHCII status, No Relapse, Early Relapse) revealed proteins and spatial metrics biomarker candidates for these clinical factors (FIG 5E, SFIG6H-J). MHCII status was strongly associated with spatial proximity to HLADPDQDR HRS tumor cells, HLADPDQDR expression, and HLADPDQDR+ HRS cells as expected (FIG 5L). Comparing spatial measurements only and their relative significance to specific cell types (FIG 5J), Macrophage and HRS spatial measurements were most consistently significant in relation to other cell types, while Endothelial, Treg, and cDC spatial measurements were rarely significant.

## DISCUSSION

This spatial protein study of matched diagnostic and relapse samples of relapsed/refractory Hodgkin Lymphoma is a resource with analysis tailored towards biomarker discovery. We identified expression patterns that appear significant to predicting outcomes in HL across different intercellular distances (FIG 6). In the unselected regions, CD163+ Macrophages^47^, GranzymeB+CD8+ T cells^39, 40^, and various HRS subtypes^30, 38, 49, 54^ were significant predictors, as previously reported. In local microenvironment niches, significant patterns were observed in CD80, TIM3, and PDL1 on macrophages, and CXCR5^83^ on HRS cells. At the hyper-local cell contact level, ligand-receptor interactions appear to drive aggregate formation, involving well known (LAG3/HLAII with CD4+ T cells, PD1/PDL1 with CD4+ and CD8+ T cells, TIM3/Galectin9 with CD8+ T cells and macrophages^27, 69, 78^) and proposed (VISTA/Galectin9 with CD4+ and CD8+ T cell^74^) signaling pairs. Alongside these promising biomarker candidates, our analysis also generates negative results. For instance, the study that proposed local expression of HLA-ABC’s usefulness as a biomarker (FIG 4D) also proposed that local expression of HLA-DPDQDR would not be significant, which we also corroborated (p=0.24).

**Figure 6.**
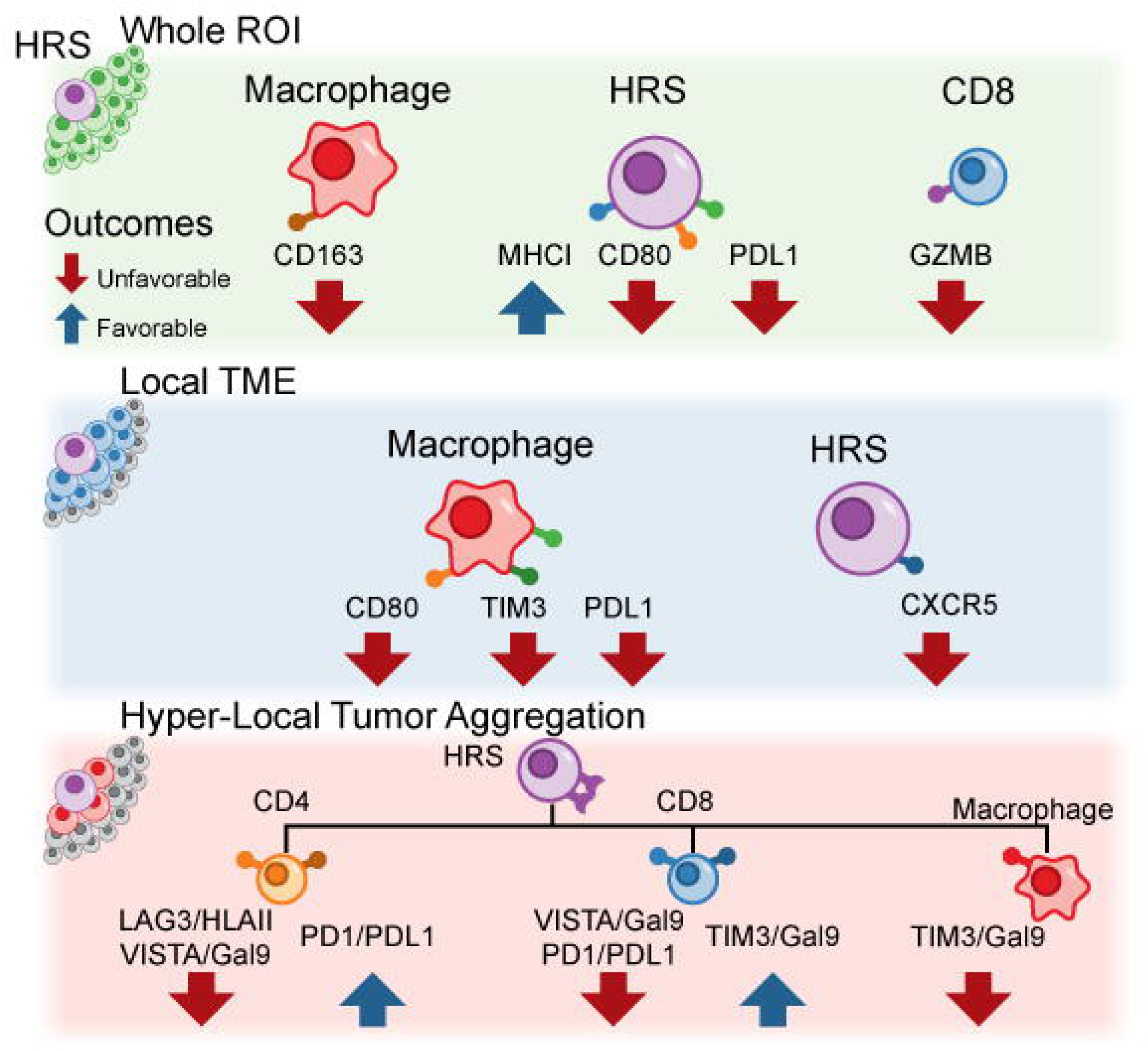
Prognostic protein expression patterns. Considering whole TMA-sized ROIs, classically recognized cell types such as CD163+ macrophages predicted early relapse and poor survival. Using spatial subsets of images via digital biopsies, additional macrophage and HRS protein expression patterns were prognostic. In ligand/receptor aggregation analysis, T cell and macrophage checkpoint patterns were significantly associated with early relapses.

We propose that the hyper-local aggregate biomarkers have 3 primary advantages. First, they incorporate both tumor and immune signaling information. Second, they are calculated within samples and may therefore be less sensitive to technical issues such as batch effects or heterogeneous tissue sampling. Third, they are spatially restricted and not overreaching. We can propose a simple hypothesis for aggregate biomarkers, where upregulated checkpoint expression between tumor cells and their immediate neighbors is more indicative of checkpoint signaling between tumor and specific immune cell types than other non-spatial measurements such as global expression of immune checkpoints or immune abundance. While larger spatial niches involving tens to hundreds of cells were also found with statistically significant survival associations in our analysis, they are more descriptive in nature and are too complex to suggest a mechanistic basis for their formation.

Spatial analytics is undergoing a modern transformation by the introduction of highly multiplexed technologies with single cell-resolution^34, 61, 84–86^. Clinical use of multiplexed IHC of up to 7 markers is becoming standardized (CLIA approved)^87–90^. Further multiplexing to 100+ marker analysis possible using cyclic imagers^61, 91, 92^. There are several distinctions between this pipeline and others^70, 93–96^. First, we used single cell spatial distance metrics to perform spatial and microenvironment analysis, instead of community-or graph-based strategies^61, 62^. We also added biomarker candidate discovery and validation tools here to address the gap between spatial IMC analysis and translation to clinical biomarkers. Unique to our study, we allowed cells to carry multiple labels of user-defined categories. Our approach was tailored towards clinical studies, which are often structured around single protein-or cell type-based hypotheses that can be lost or minimized by dimensional reduction. This design interfaced with our hypothesis testing and biomarker candidate validation tools, which guide the user towards the generation of standardized Kaplan-Meier survival curves and hazard ratios as well as spatially-informed biomarkers.

We proposed IMC as a tissue-efficient strategy for biomarker discovery and validation. We emphasized 4 previous biomarker studies (Fig 4A-D, CD68+ macrophages, GranzymeB+CD8+ T cells, PDL1+ HRS, MHCI-HRS, FIG 4A-D^24, 30, 39, 49^) that collectively represented 634 patients. Each study represents a significant investment in time and resources to validate, which was condensed into this single study using IMC. Our spatial strategy using digital biopsies also mitigated the problem of tumor heterogeneity causing biases in IMC-sized ROIs sampled from tissue. This strategy applies to all imaging studies, which will be important until the price of analyzing whole slides decreases. IMC can be applied broadly to identify parsimonious combinations of 3-5 proteins as biomarkers for clinical grade multi-color IHC platforms^89, 90, 97^. Similar biomarkers are already mainstays in routine diagnostic procedures for subtyping with implications for clinical prognosis, for example the Hans algorithm in diffuse large B cell lymphoma that was developed decades ago (Hans, etc)^98, 99^.

While the use of such highly multiplexed imaging as used in this study is not currently feasible in routine pathology diagnostics, findings from this study offer novel insights into prognostic elements based on spatial patterns. Among the biomarker candidates generated by our analysis, we identified candidate proteins such as PDL1, TIM3, LAG3, and CXCR5 deserving of further attention and identified the specific cell types and contexts for potential biomarkers. For HL, the relatively high success rate of ABVD chemotherapy suggests that secondary outcomes, such as 1^st^ or 2^nd^ Failure Free Survival (FFS) and postBMT FFS, may be more clinically significant. We performed a companion study to this one focused on translating IMC studies to a biomarker assay for postBMT FFS using IHC^100^. In that study, we presented additional strategies to refine LASSO_plus variable selection, and demonstrated a protocol to select a parsimonious biomarker set of 6 proteins for IHC and standardize IMC and IHC data comparisons. We used the protocol to validate a new HL biomarker (RHL4S), based on the spatial arrangement of CXCR5+ HRS cells, PD1+CD4+ T cells, tumor-associated macrophages, and CXCR5+ B cells. With the coming of age of digital pathology and incorporation of multiplexed IHC into routine diagnostics, methods to identify these spatial patterns will become increasingly important for effective prognostication.

Spatial context expression requires interpretation, as in the aggregate studies we presented here. For instance, HRS cell checkpoint expression counterintuitively increased with CD4+ T cell aggregate size, especially in late relapses. Such HRS cells may lack other immunosuppressive mechanisms which then allows immune cells to be recruited to the HRS aggregates despite checkpoint expression. Non relapse, late relapse, and early relapse disease are also too complex to fall neatly in a continuum. One hypothesis is that late relapses, which occurred as late as 18 years after diagnosis, are due to failures in immune surveillance, which may manifest in our observed differences in aggregate composition and protein expression that are unique to late relapse patients, such as the increased frequency of CD8+ T cell aggregates (FIG 4F-G, SFIG5B). By systematically quantifying local aggregate signaling we can validate spatial signaling insights predicted by other methods^101–104^.

Cell segmentation of spatial data remains challenging especially in large cells or cells with concave or serpentine shapes in tissue (HRS, DCs), despite the recent application of machine learning in tools that require little to no user input (Mesmer^105^) as opposed to the supervised tools used here (ilastik^106^). With imperfect segmentation and ambiguous cell phenotyping, human input is used to simplify data, usually by omitting problematic cells^34^, or binarizing or simplifying the expression of phenotyping markers to remove ambiguity^107, 108^. In our study, we found that HRS-ICOS/CD4-ICOSL was observed as a highly significant association with CD4+ T cell aggregate size. However, ICOS is not believed to be expressed on HRS cells. Thus, “expression” of ICOS may be a technical observation associated with increased clustering of CD4+ T cells and imperfect segmentation, rather than biological HRS cell expression, which is reinforced by the universal significance of this protein expression pattern across all timepoints and patient outcomes.

Commercial spatial transcriptomics^109–113^ now present an alternative spatial tool to spatial protein analysis, and the strengths and weaknesses of the technologies must be weighed. Currently, spatial transcriptomics is limited to the discovery setting due to its cost, and we believe the translational biomarker pipeline established here represents an essential use case for IMC. The amount of biomarker candidates generated by IMC is beyond what can be reasonably tested and validated using separate gold standard multi-color IHC experiments, the materials and reagents used in IMC can be rapidly translated to IHC in clinical settings, and the tools presented here can accelerate translational biomarker studies in a variety of diseases and clinical challenges.

## METHODS

### SAMPLE COLLECTION AND TMA ASSEMBLY DETAILS

The patient cohort was described in a previous publication, which analyzed gene expression profiling of relapsed and refractory HL^114^. We analyzed a tissue microarray of a cohort of 93 total HL patients treated at British Columbia Cancer comprising 73 diagnostic and relapsed paired samples and 20 primary biopsies of patients who were cured after ABVD treatment for a total of 256 ROIs with up to 2 ROIs per biopsies when available (average 1.76). Patients were selected according to the following criteria: patients received first-line treatment with doxorubicin, bleomycin, vinblastine, and dacarbazine (ABVD) or ABVD-equivalent therapy with curative intent; patients experienced CHL progression/relapse after primary (refractory disease or relapse); and tissue derived from an excisional biopsy was available^33^. This study was reviewed and approved by the University of British Columbia-BC Cancer Agency Research Ethics Board (H14-02304), in accordance with the Declaration of Helsinki. We obtained written informed consent from the patients or informed consent was waived for the samples used in this retrospective study.

### ANTIBODY CONJUGATIONS

Antibodies were purchased from Fluidigm Inc., now Standard Biotools, in conjugated form. Antibodies that were unavailable were conjugated using MaxPar kits. TMA slides were dewaxed in 3 washes of xylene and rehydrated by successive washes in 100%, 95%, 80%, and 70% ethanol in distilled water. After washing with the alcohol gradient, the slides were immersed in Tris-EDTA antigen retrieval solution for 30 minutes at 95°C and were left to cool down inside the solution for 30 minutes more at room temperature. After the antigen retrieval step the slides were blocked with 3% BSA for 45 minutes and then were stained overnight with the antibody panel at 4°C. The next day the slides were washed twice with PBS-0.1% Triton X solution and 1X PBS for 8 minutes each. The slides were then incubated with 191 Iridium, a nuclear stain, for 40 minutes. After that, the slides were washed with dH_2_O and dried off before ablation. The slides were ablated using the Hyperion/Helios Imaging Mass Cytometry platform (Standard Biotools) at a rate of 200 H, and images were acquired for analysis.

### IMAGE ANALYSIS

Raw image files were generated using MCD Viewer (Standard Biotools). Segmentation was performed using the IMC Segmentation Pipeline (Bodenmiller lab) with modification. First, a subset of antibodies was chosen for Ilastik training. We define groups of membrane-specific (CD14, HLA-ABC, CD68, CD31, CD4, CD20, CD8a, CD30, CD3, CD45RO, HLA-DPDQDR) and nuclear-specific markers (Ir191, Ir193) and add 4 additional channels to the Ilastik training: the sum of all antibody channels, the sum of nuclear-specific channels, the sum-of membrane-specific channels, and a Sobel-filtered nuclear channel which highlights the edges of nuclei by emphasizing sharp changes and de-emphasizing constant regions of nuclear signal. We did not quantify the effect of adding these channels as each Ilastik training is a highly specific and unique series of drawn regions for every image.

Segmented cell data was processed using Python and R. Cells smaller than 5 μm^2^ or larger than 300 μm^2^ were removed from analysis, and ROIs with fewer than 500 cells were removed from analysis. Particular challenges in segmentation of IMC data included some multi-nucleated cells, as well as cell fragments due to the nature of tissue slices. HRS and DC cells were likely most affected by these limitations. Mean cell protein values were transformed by hyperbolic arcsin, scaled, censored at the top 1%, and all clustering steps were performed using Rphenograph with k=15. Automated clustering was performed by calculating median protein values of protein expression, normalizing to z-scores, and gating clusters. Cell type phenotype labeling and triage for reclustering was performed manually at the cluster level. Cells were allowed to carry multiple phenotype labels.

To perform quality control on phenotyping, we measure the heterogeneity of cell prevalence across different measurement units (i.e. patients) using the Shannon informational entropy (SFIG2B). For any cell type, entropy is minimized when all cells are found in a single patient, and it is maximized when every patient has the same proportion of that cell type. While we do not expect every cell to be found at the same frequency in each patient (maximum entropy), very low entropy can be indicative of batch-or patient-specific-effects.

Our script presents two strategies for phenotyping depending on the size of the data set: full clustering (SFIG2A) or metaclustering. Beginning with the first tier of cell phenotyping markers, full clustering uses a single clustering step for every cell, which can be computationally intensive as data sets grow beyond 1 million cells and more than 10 proteins are used. Metaclustering (used in this study) first performs clustering at a smaller subset level (image-level, patient-level), then performs clustering of mean expression levels of clusters. Automatic labels were assigned by calculating z-scores of protein expression for clusters and labeling each cluster with all cell types based on protein expression.

Within each of these automatically labeled clusters, we performed a second Phenograph step and manually sorted each cluster into the cell type of interest (SFIG2D). Remaining cells unable to be classified were subjected to a final clustering step and z-scores of cell expression were used to assign cell types, with multiple cell type assignments per cell allowed. After manual curation, each major phenotype underwent a Phenograph step using cell-type-specific markers and relevant phenotype subtypes of interest were defined. UMAPs were generated using the uwot package for all cells as well as cell subtypes using default settings.

### SPATIAL ANALYSIS

For spatial analysis, the distance of each cell to its 5 nearest neighbors of every cell phenotype label was calculated, capped at 50 μm, scaled from 0-1, and inverted. These distance metrics describe the local enrichment for a specific cell type to each cell, and were used in clustering using k-means clustering (k=15 or 35) too define spatial niches/local microenvironments. Local aggregation-dependent protein expression was obtained by selecting aggregating cell types (CD4+ T, CD8+ T, Treg, macrophage, B cell), and identifying all cells of the aggregating cell type within the contact radius (15 μm) of every cell. The number of aggregating cells within the radius and the mean protein expression were recorded. Ligand receptor expression was defined as the product of the ligand and receptor on the central cell and aggregating cell average, respectively. Aggregate-specific protein expression was calculated by constructing a generalized linear model of aggregate size as a function of aggregating cell protein, HRS cell protein, or ligand-receptor expression, and extracting p values for each of the biomarker candidates.

### AGGREGATE SUBSTITUTION TEST

Each ROI was independently analyzed for random replacement testing of aggregation frequency. For each type of aggregating cell, phenotype labels were reshuffled on every cell to match the original cell proportions, and the number of aggregates around HRS cells was calculated (defined by at least 3 aggregating cells). This process was performed 10,000 times, and the proportion of times for which the reshuffled image contained more aggregates than the observed aggregates was defined as the test fraction.

### BIOMARKER ANALYSIS

For phenotype biomarkers, each ROI was summarized by the percent composition of every labeled phenotype. We used the mclust R package at its default settings to perform model-based clustering to stratify patients. Stratifications that produced greater than a 90:10 split were removed from analysis. Cox proportional hazards survival analysis was performed using the survival package in R. Biomarker analysis was performed on subsets of the data separated by temporal conditions (diagnostic/relapse, 2 conditions) and spatial conditions (whole image, tumor enrichment – cells with HRS enrichment > 0.5, and tumor contact – cells with HRS nearest neighbor distance < 10 μm), which we defined as “digital biopsies” (3 conditions). A total of 5 possible event codes (overall survival, disease-specific survival, freedom from first failure survival, freedom from second failure survival, and bone marrow transplant freedom from failure survival) were available^33^, resulting in 30 possible survival analyses across the temporal and spatial conditions. Each biomarker candidate that was significant in at least 4 conditions was retained, and p-values were adjusted by the Benjamini-Hochberg false discovery rate correction within these retained candidates.

A linear or generalized linear model was used to analyze single-cell IMC data for continuous or categorical outcomes. The sample was considered as a clustering variable, and cluster-robust standard errors were computed. The analysis was conducted using the ‘stats’ package in R. Cox regression is readily used with patient-level biomarkers, and for single-cell biomarkers, Cox regression can be performed with a cluster or frailty variable, mixed models and generalized estimating equations, or by downsampling each patient to obtain equal numbers of cells to ensure that each patient is weighted properly. These strategies have not performed well in the IMC setting, and we recommend converting cell-level biomarkers into patient-level biomarkers to perform Cox regression as done here using the mean.

### LASSO_plus

Using sample-level data, we present a pipeline using LASSO_plus for variable selection, which is available in the R package “csmpv” (https://github.com/ajiangsfu/csmpv). This variable selection pipeline is processed with three steps using either cell proportions, mean cell-type-specific protein expression, or mean-cell-type-specific spatial metrics as variables. The 1^st^ step is a customized version of LASSO. Variable selection is performed with the glmnet R package to extract candidate biomarkers and variables that provide minimal information are penalized and removed^115, 116^. In our customized LASSO step, a pre-defined top N (default setting is 10) is used to select a stable list of variables instead of LASSO’s variable selection method. To do that, we record numbers of variables for all lambda simulations, keep only the repeated numbers, then select the repeated number that is closest to the pre-defined top N.

The 2^nd^ step is single variable selection which aims to salvage any variables that were excluded by LASSO as they shared redundant information with the already selected variables. This is achieved using single independent variable regression, which can be performed using either a linear model, a generalized linear model, or a Cox model, depending on the type of outcome being studied. The 3^rd^ step is stepwise model selection using the combined list of variables obtained from LASSO and single variable selection^117^. The criterion for model comparison is the Akaike Information Criterion (AIC). The approach entails gradually building up a model by including or excluding one variable at a time. Ultimately, the model with the lowest AIC score among all potential models is chosen as the best model, and the variables in this model make up the final selected variable list.

## Notes

https://zenodo.org/deposit/7963681

### Competing Interest Statement

C.S. has performed consultancy for Bayer and has received research funding from Epizyme and Trillium Therapeutics. C.S. is a named inventor on a patent filed by the National Cancer Institute Methods for determining lymphoma type. The remaining authors declare no competing financial interests.

### Summary of Updates

Minor edit to figure 3, small image was missing.

## REFERENCES

1. C. Randall and Y. Fedoriw, Pathology, 2020, 52, 30–39.

2. K. H. Allison, American journal of clinical pathology, 2012, 138, 770–780.

3. W. D. Travis, Clin Chest Med, 2011, 32, 669–692.

4. A. H. Coons, H. J. Creech, R. N. Jones and E. Berliner, The Journal of Immunology, 1942, 45, 159–170.

5. A. H. Fischer, K. A. Jacobson, J. Rose and R. Zeller, CSH Protoc, 2008, 2008, pdb prot4986.

6. S. W. Kim, J. Roh and C. S. Park, J Pathol Transl Med, 2016, 50, 411–418.

7. C. E. Cox, S. Pendas, J. M. Cox, E. Joseph, A. R. Shons, T. Yeatman, N. N. Ku, G. H. Lyman, C. Berman, F. Haddad and D. S. Reintgen, Ann Surg, 1998, 227, 645–651; discussion 651-643.

8. I. Anagnostopoulos, M. L. Hansmann, K. Franssila, M. Harris, N. L. Harris, E. S. Jaffe, J. Han, J. M. van Krieken, S. Poppema, T. Marafioti, J. Franklin, M. Sextro, V. Diehl and H. Stein, Blood, 2000, 96, 1889–1899.

9. S. Hartmann, S. Scharf, Y. Steiner, A. G. Loth, E. Donnadieu, N. Flinner, V. Poeschel, S. Angel, M. Bewarder, J. Bein, U. Brunnberg, A. Bozzato, B. Schick, S. Stilgenbauer, R. M. Bohle, L. Thurner and M. L. Hansmann, Cancers (Basel), 2021, 13.

10. R. Alaggio, C. Amador, I. Anagnostopoulos, A. D. Attygalle, I. B. O. Araujo, E. Berti, G. Bhagat, A. M. Borges, D. Boyer, M. Calaminici, A. Chadburn, J. K. C. Chan, W. Cheuk, W. J. Chng, J. K. Choi, S. S. Chuang, S. E. Coupland, M. Czader, S. S. Dave, D. de Jong, M. Q. Du, K. S. Elenitoba-Johnson, J. Ferry, J. Geyer, D. Gratzinger, J. Guitart, S. Gujral, M. Harris, C. J. Harrison, S. Hartmann, A. Hochhaus, P. M. Jansen, K. Karube, W. Kempf, J. Khoury, H. Kimura, W. Klapper, A. E. Kovach, S. Kumar, A. J. Lazar, S. Lazzi, L. Leoncini, N. Leung, V. Leventaki, X. Q. Li, M. S. Lim, W. P. Liu, A. Louissaint, Jr., A. Marcogliese, L. J. Medeiros, M. Michal, R. N. Miranda, C. Mitteldorf, S. Montes-Moreno, W. Morice, V. Nardi, K. N. Naresh, Y. Natkunam, S. B. Ng, I. Oschlies, G. Ott, M. Parrens, M. Pulitzer, S. V. Rajkumar, A. C. Rawstron, K. Rech, A. Rosenwald, J. Said, C. Sarkozy, S. Sayed, C. Saygin, A. Schuh, W. Sewell, R. Siebert, A. R. Sohani, R. Tooze, A. Traverse-Glehen, F. Vega, B. Vergier, A. D. Wechalekar, B. Wood, L. Xerri and W. Xiao, Leukemia, 2022, 36, 1720–1748.

11. E. Campo, E. S. Jaffe, J. R. Cook, L. Quintanilla-Martinez, S. H. Swerdlow, K. C. Anderson, P. Brousset, L. Cerroni, L. de Leval, S. Dirnhofer, A. Dogan, A. L. Feldman, F. Fend, J. W. Friedberg, P. Gaulard, P. Ghia, S. M. Horwitz, R. L. King, G. Salles, J. San-Miguel, J. F. Seymour, S. P. Treon, J. M. Vose, E. Zucca, R. Advani, S. Ansell, W. Y. Au, C. Barrionuevo, L. Bergsagel, W. C. Chan, J. I. Cohen, F. d’Amore, A. Davies, B. Falini, I. M. Ghobrial, J. R. Goodlad, J. G. Gribben, E. D. Hsi, B. S. Kahl, W. S. Kim, S. Kumar, A. S. LaCasce, C. Laurent, G. Lenz, J. P. Leonard, M. P. Link, A. Lopez-Guillermo, M. V. Mateos, E. Macintyre, A. M. Melnick, F. Morschhauser, S. Nakamura, M. Narbaitz, A. Pavlovsky, S. A. Pileri, M. Piris, B. Pro, V. Rajkumar, S. T. Rosen, B. Sander, L. Sehn, M. A. Shipp, S. M. Smith, L. M. Staudt, C. Thieblemont, T. Tousseyn, W. H. Wilson, T. Yoshino, P. L. Zinzani, M. Dreyling, D. W. Scott, J. N. Winter and A. D. Zelenetz, Blood, 2022, 140, 1229–1253.

12. D. Hasenclever and V. Diehl, The New England journal of medicine, 1998, 339, 1506–1514.

13. A. A. Moccia, J. Donaldson, M. Chhanabhai, P. J. Hoskins, R. J. Klasa, K. J. Savage, T. N. Shenkier, G. W. Slack, B. Skinnider, R. D. Gascoyne, J. M. Connors and L. H. Sehn, Journal of Clinical Oncology, 2012, 30, 3383–3388.

14. A. Gallamini, M. Hutchings, L. Rigacci, L. Specht, F. Merli, M. Hansen, C. Patti, A. Loft, F. Di Raimondo, F. D’Amore, A. Biggi, U. Vitolo, C. Stelitano, R. Sancetta, L. Trentin, S. Luminari, E. Iannitto, S. Viviani, I. Pierri and A. Levis, Journal of clinical oncology: official journal of the American Society of Clinical Oncology, 2007, 25, 3746–3752.

15. J. Radford, T. Illidge, N. Counsell, B. Hancock, R. Pettengell, P. Johnson, J. Wimperis, D. Culligan, B. Popova, P. Smith, A. McMillan, A. Brownell, A. Kruger, A. Lister, P. Hoskin, M. O’Doherty and S. Barrington, New England Journal of Medicine, 2015, 372, 1598–1607.

16. P. Johnson, M. Federico, A. Kirkwood, A. Fosså, L. Berkahn, A. Carella, F. d’Amore, G. Enblad, A. Franceschetto, M. Fulham, S. Luminari, M. O’Doherty, P. Patrick, T. Roberts, G. Sidra, L. Stevens, P. Smith, J. Trotman, Z. Viney, J. Radford and S. Barrington, The New England journal of medicine, 2016, 374, 2419–2429.

17. S. Shanbhag and R. F. Ambinder, CA Cancer J Clin, 2018, 68, 116–132.

18. J. M. Connors, W. Jurczak, D. J. Straus, S. M. Ansell, W. S. Kim, A. Gallamini, A. Younes, S. Alekseev, Á. Illés, M. Picardi, E. Lech-Maranda, Y. Oki, T. Feldman, P. Smolewski, K. J. Savage, N. L. Bartlett, J. Walewski, R. Chen, R. Ramchandren, P. L. Zinzani, D. Cunningham, A. Rosta, N. C. Josephson, E. Song, J. Sachs, R. Liu, H. A. Jolin, D. Huebner and J. Radford, New England Journal of Medicine, 2017, 378, 331–344.

19. A. Younes, A. K. Gopal, S. E. Smith, S. M. Ansell, J. D. Rosenblatt, K. J. Savage, R. Ramchandren, N. L. Bartlett, B. D. Cheson, S. de Vos, A. Forero-Torres, C. H. Moskowitz, J. M. Connors, A. Engert, E. K. Larsen, D. A. Kennedy, E. L. Sievers and R. Chen, Journal of clinical oncology: official journal of the American Society of Clinical Oncology, 2012, 30, 2183–2189.

20. C. H. Moskowitz, A. Nademanee, T. Masszi, E. Agura, J. Holowiecki, M. H. Abidi, A. I. Chen, P. Stiff, A. M. Gianni, A. Carella, D. Osmanov, V. Bachanova, J. Sweetenham, A. Sureda, D. Huebner, E. L. Sievers, A. Chi, E. K. Larsen, N. N. Hunder and J. Walewski, The Lancet, 2015, 385, 1853–1862.

21. G. P. Canellos, J. R. Anderson, K. J. Propert, N. Nissen, M. R. Cooper, E. S. Henderson, M. R. Green, A. Gottlieb and B. A. Peterson, New England Journal of Medicine, 1992, 327, 1478–1484.

22. L. I. Gordon, F. Hong, R. I. Fisher, N. L. Bartlett, J. M. Connors, R. D. Gascoyne, H. Wagner, P. J. Stiff, B. D. Cheson, M. Gospodarowicz, R. Advani, B. S. Kahl, J. W. Friedberg, K. A. Blum, T. M. Habermann, J. M. Tuscano, R. T. Hoppe and S. J. Horning, Journal of Clinical Oncology, 2012, 31, 684–691.

23. T. W. Kelley, B. Pohlman, P. Elson and E. D. Hsi, American journal of clinical pathology, 2007, 128, 958–965.

24. C. Steidl, T. Lee, S. P. Shah, P. Farinha, G. Han, T. Nayar, A. Delaney, S. J. Jones, J. Iqbal, D. D. Weisenburger, M. A. Bast, A. Rosenwald, H. K. Muller-Hermelink, L. M. Rimsza, E. Campo, J. Delabie, R. M. Braziel, J. R. Cook, R. R. Tubbs, E. S. Jaffe, G. Lenz, J. M. Connors, L. M. Staudt, W. C. Chan and R. D. Gascoyne, The New England journal of medicine, 2010, 362, 875–885.

25. K. Jones, F. Vari, C. Keane, P. Crooks, J. P. Nourse, L. A. Seymour, D. Gottlieb, D. Ritchie, D. Gill and M. K. Gandhi, Clin Cancer Res, 2013, 19, 731–742.

26. T. Aoki and C. Steidl, Cancer J, 2018, 24, 206–214.

27. C. D. Carey, D. Gusenleitner, M. Lipschitz, M. G. M. Roemer, E. C. Stack, E. Gjini, X. Hu, R. Redd, G. J. Freeman, D. Neuberg, F. S. Hodi, X. S. Liu, M. A. Shipp and S. J. Rodig, Blood, 2017, 130, 2420–2430.

28. M. Kawashima, J. Carreras, H. Higuchi, R. Kotaki, T. Hoshina, K. Okuyama, N. Suzuki, M. Kakizaki, Y. Miyatake, K. Ando, M. Nakayama, S. Umezu, R. Horie, Y. Higuchi, K. Katagiri, S. Goyama, T. Kitamura, K. Chamoto, S. Yano, N. Nakamura and A. Kotani, Leukemia, 2020, 34, 2405–2417.

29. S. S. Patel, J. L. Weirather, M. Lipschitz, A. Lako, P.-H. Chen, G. K. Griffin, P. Armand, M. A. Shipp and S. J. Rodig, Blood, 2019, 134, 2059–2069.

30. M. G. Roemer, R. H. Advani, R. A. Redd, G. S. Pinkus, Y. Natkunam, A. H. Ligon, C. F. Connelly, C. J. Pak, C. D. Carey, S. E. Daadi, B. Chapuy, D. de Jong, R. T. Hoppe, D. S. Neuberg, M. A. Shipp and S. J. Rodig, Cancer Immunol Res, 2016, 4, 910–916.

31. J. Veldman, L. Visser, M. Huberts-Kregel, N. Muller, B. Hepkema, A. van den Berg and A. Diepstra, Blood, 2020, 136, 2437–2441.

32. D. J. Flavell and D. H. Wright, Br J Cancer, 1989, 59, 165–173.

33. F. C. Chan, A. Mottok, A. S. Gerrie, M. Power, M. Nijland and A. Diepstra, Journal of clinical oncology: official journal of the American Society of Clinical Oncology, 2017, 35, 3722–3733.

34. C. Giesen, H. A. Wang, D. Schapiro, N. Zivanovic, A. Jacobs, B. Hattendorf, P. J. Schuffler, D. Grolimund, J. M. Buhmann, S. Brandt, Z. Varga, P. J. Wild, D. Gunther and B. Bodenmiller, Nat Methods, 2014, 11, 417–422.

35. H. W. Jackson, J. R. Fischer, V. R. T. Zanotelli, H. R. Ali, R. Mechera, S. D. Soysal, H. Moch, S. Muenst, Z. Varga, W. P. Weber and B. Bodenmiller, Nature, 2020, 578, 615–620.

36. D. Schulz, V. R. T. Zanotelli, J. R. Fischer, D. Schapiro, S. Engler, X. K. Lun, H. W. Jackson and B. Bodenmiller, Cell Syst, 2018, 6, 25–36 e25.

37. U. Schwab, H. Stein, J. Gerdes, H. Lemke, H. Kirchner, M. Schaadt and V. Diehl, Nature, 1982, 299, 65–67.

38. M. M. Morente, M. A. Piris, V. Abraira, A. Acevedo, B. Aguilera, C. Bellas, M. Fraga, R. Garcia-Del-Moral, F. Gomez-Marcos, J. Menarguez, H. Oliva, M. Sanchez-Beato and C. Montalban, Blood, 1997, 90, 2429–2436.

39. J. J. Oudejans, N. M. Jiwa, J. A. Kummer, G. J. Ossenkoppele, P. van Heerde, J. W. Baars, P. M. Kluin, J. C. Kluin-Nelemans, P. J. van Diest, J. M. Middeldorp and C. J. L. M. Meijer, Blood, 1997, 89, 1376–1382.

40. R. L. ten Berge, J. J. Oudejans, D. F. Dukers, J. W. R. Meijer, G. J. Ossenkoppele and C. Meijer, Leukemia, 2001, 15, 458–464.

41. D. Aldinucci, D. Poletto, A. Gloghini, P. Nanni, M. Degan, T. Perin, P. Ceolin, F. M. Rossi, V. Gattei, A. Carbone and A. Pinto, The American journal of pathology, 2002, 160, 585–596.

42. A. Tzankov, J. Krugmann, F. Fend, M. Fischhofer, R. Greil and S. Dirnhofer, Clin Cancer Res, 2003, 9, 1381–1386.

43. T. s. Álvaro, M. n. Lejeune, M. T. Salvadó, R. n. Bosch, J. F. García, J. n. Jaén, A. H. Banham, G. Roncador, C. Montalbán and M. A. Piris, Clinical Cancer Research, 2005, 11, 1467–1473.

44. S. Muenst, S. Hoeller, S. Dirnhofer and A. Tzankov, Human pathology, 2009, 40, 1715–1722.

45. S. Schreck, D. Friebel, M. Buettner, L. Distel, G. Grabenbauer, L. S. Young and G. Niedobitek, Hematol Oncol, 2009, 27, 31–39.

46. M. H. M. Barros, R. Hassan and G. Niedobitek, Clinical Cancer Research, 2012, 18, 3762–3771.

47. K. L. Tan, D. W. Scott, F. Hong, B. S. Kahl, R. I. Fisher, N. L. Bartlett, R. H. Advani, R. Buckstein, L. M. Rimsza, J. M. Connors, C. Steidl, L. I. Gordon, S. J. Horning and R. D. Gascoyne, Blood, 2012, 120, 3280–3287.

48. P. Greaves, A. Clear, R. Coutinho, A. Wilson, J. Matthews, A. Owen, M. Shanyinde, T. A. Lister, M. Calaminici and J. G. Gribben, Journal of clinical oncology: official journal of the American Society of Clinical Oncology, 2013, 31, 256–262.

49. T. Menter, A. Bodmer-Haecki, S. Dirnhofer and A. Tzankov, Human pathology, 2016, 54, 17–24.

50. R. Sun, L. J. Medeiros and K. H. Young, Modern Pathology, 2016, 29, 1118–1142.

51. F. Z. Cader, R. C. J. Schackmann, X. Hu, K. Wienand, R. Redd, B. Chapuy, J. Ouyang, N. Paul, E. Gjini, M. Lipschitz, P. Armand, D. Wu, J. R. Fromm, D. Neuberg, X. S. Liu, S. J. Rodig and M. A. Shipp, Blood, 2018, 132, 825–836.

52. S. Péricart, M. Tosolini, P. Gravelle, C. Rossi, A. Traverse-Glehen, N. Amara, C. Franchet, E. Martin, C. Bezombes, G. Laurent, P. Brousset, J.-J. Fournié and C. Laurent, Cancers, 2018, 10, 415.

53. M. G. M. Roemer, R. A. Redd, F. Z. Cader, C. J. Pak, S. Abdelrahman, J. Ouyang, S. Sasse, A. Younes, M. Fanale, A. Santoro, P. L. Zinzani, J. Timmerman, G. P. Collins, R. Ramchandren, J. B. Cohen, J. P. De Boer, J. Kuruvilla, K. J. Savage, M. Trneny, S. Ansell, K. Kato, B. Farsaci, A. Sumbul, P. Armand, D. S. Neuberg, G. S. Pinkus, A. H. Ligon, S. J. Rodig and M. A. Shipp, Journal of clinical oncology: official journal of the American Society of Clinical Oncology, 2018, 36, 942–950.

54. S. Jalali, T. Price-Troska, C. Bothun, J. Villasboas, H.-J. Kim, Z.-Z. Yang, A. J. Novak, H. Dong and S. M. Ansell, Blood Cancer Journal, 2019, 9, 22.

55. K. Karihtala, S. K. Leivonen, O. Brück, M. L. Karjalainen-Lindsberg, S. Mustjoki, T. Pellinen and S. Leppä, Cancers (Basel), 2020, 12.

56. S. Moerdler, M. Ewart, D. L. Friedman, K. Kelly, Q. Pei, M. Peng, X. Zang and P. D. Cole, Leuk Lymphoma, 2021, 62, 606–613.

57. A. Santisteban-Espejo, I. Bernal-Florindo, J. Perez-Requena, L. Atienza-Cuevas, N. Maira-Gonzalez and M. Garcia-Rojo, Frontiers in Oncology, 2022, 12.

58. M. A. Piris, L. J. Medeiros and K. C. Chang, Pathology, 2020, 52, 154–165.

59. P. J. Clark and F. C. Evans, Ecology, 1954, 35, 445–453.

60. E. Calabretta, F. d’Amore and C. Carlo-Stella, Int J Mol Sci, 2019, 20.

61. C. M. Schurch, S. S. Bhate, G. L. Barlow, D. J. Phillips, L. Noti, I. Zlobec, P. Chu, S. Black, J. Demeter, D. R. McIlwain, S. Kinoshita, N. Samusik, Y. Goltsev and G. P. Nolan, Cell, 2020, 182, 1341–1359 e1319.

62. K. H. Gouin, 3rd, N. Ing, J. T. Plummer, C. J. Rosser, B. Ben Cheikh, C. Oh, S. S. Chen, K. S. Chan, H. Furuya, W. G. Tourtellotte, S. R. V. Knott and D. Theodorescu, Nat Commun, 2021, 12, 4906.

63. D. Voduc, C. Kenney and T. O. Nielsen, Semin Radiat Oncol, 2008, 18, 89–97.

64. A. Hoos and C. Cordon-Cardo, Laboratory Investigation, 2001, 81, 1331–1338.

65. S. M. Hewitt, Methods Mol Biol, 2012, 823, 201–214.

66. E. A. G. Baker, D. Schapiro, B. Dumitrascu, S. Vickovic and A. Regev, bioRxiv, 2022, DOI: 10.1101/2022.01.26.477748, 2022.2001.2026.477748.

67. J. MacQueen, 1967.

68. R. L. Thorndike, Psychometrika, 1953, 18, 267–276.

69. L. El Halabi, J. Adam, P. Gravelle, V. Marty, A. Danu, J. Lazarovici, V. Ribrag, J. Bosq, V. Camara-Clayette, C. Laurent and D. Ghez, Clinical Lymphoma Myeloma and Leukemia, 2021, 21, 257–266.e253.

70. D. Schapiro, H. W. Jackson, S. Raghuraman, J. R. Fischer, V. R. T. Zanotelli, D. Schulz, C. Giesen, R. Catena, Z. Varga and B. Bodenmiller, Nat Methods, 2017, 14, 873–876.

71. D. R. Leach, M. F. Krummel and J. P. Allison, Science, 1996, 271, 1734–1736.

72. J. M. Kim and D. S. Chen, Ann Oncol, 2016, 27, 1492–1504.

73. T. Wu and Y. Dai, Cancer Lett, 2017, 387, 61–68.

74. I. M. Yasinska, N. H. Meyer, S. Schlichtner, R. Hussain, G. Siligardi, M. Casely-Hayford, W. Fiedler, J. Wellbrock, C. Desmet, L. Calzolai, L. Varani, S. M. Berger, U. Raap, B. F. Gibbs, E. Fasler-Kan and V. V. Sumbayev, Front Immunol, 2020, 11, 580557.

75. R. Yang, L. Sun, C. F. Li, Y. H. Wang, J. Yao, H. Li, M. Yan, W. C. Chang, J. M. Hsu, J. H. Cha, J. L. Hsu, C. W. Chou, X. Sun, Y. Deng, C. K. Chou, D. Yu and M. C. Hung, Nat Commun, 2021, 12, 832.

76. C. Solinas, C. Gu-Trantien and K. Willard-Gallo, ESMO Open, 2020, 5.

77. P. Greaves, A. Clear, A. Owen, S. Iqbal, A. Lee, J. Matthews, A. Wilson, M. Calaminici and J. G. Gribben, Blood, 2013, 122, 2856–2863.

78. M. K. Gandhi, E. Lambley, J. Duraiswamy, U. Dua, C. Smith, S. Elliott, D. Gill, P. Marlton, J. Seymour and R. Khanna, Blood, 2006, 108, 2280–2289.

79. A. Sureda, M. André, P. Borchmann, M. G. da Silva, C. Gisselbrecht, T. P. Vassilakopoulos, P. L. Zinzani and J. Walewski, BMC Cancer, 2020, 20, 1088.

80. W. C. Vooijs, H. G. Otten, M. van Vliet, A. J. G. van Dijk, R. A. de Weger, M. de Boer, H. Bohlen, A. Bolognesi, L. Polito and G. C. de Gast, British Journal of Cancer, 1997, 76, 1163–1169.

81. M. Ruella, M. Klichinsky, S. S. Kenderian, O. Shestova, A. Ziober, D. O. Kraft, M. Feldman, M. A. Wasik, C. H. June and S. Gill, Cancer Discov, 2017, 7, 1154–1167.

82. T. Aoki, L. C. Chong, K. Takata, K. Milne, A. Marshall, E. A. Chavez, T. Miyata-Takata, S. Ben-Neriah, D. Unrau, A. Telenius, M. Boyle, A. P. Weng, K. J. Savage, D. W. Scott, P. Farinha, S. P. Shah, B. H. Nelson and C. Steidl, Proceedings of the National Academy of Sciences, 2021, 118, e2105822118.

83. T. Aoki, L. C. Chong, K. Takata, K. Milne, A. Marshall, E. A. Chavez, T. Miyata-Takata, S. Ben-Neriah, D. Unrau, A. Telenius, M. Boyle, A. P. Weng, K. J. Savage, D. W. Scott, P. Farinha, S. P. Shah, B. H. Nelson and C. Steidl, Proc Natl Acad Sci U S A, 2021, 118.

84. L. Keren, M. Bosse, D. Marquez, R. Angoshtari, S. Jain, S. Varma, S. R. Yang, A. Kurian, D. Van Valen, R. West, S. C. Bendall and M. Angelo, Cell, 2018, 174, 1373–1387 e1319.

85. Y. Goltsev, N. Samusik, J. Kennedy-Darling, S. Bhate, M. Hale, G. Vazquez, S. Black and G. P. Nolan, Cell, 2018, 174, 968–981.e915.

86. J. R. Lin, M. Fallahi-Sichani and P. K. Sorger, Nat Commun, 2015, 6, 8390.

87. J. M. Taube, K. Roman, E. L. Engle, C. Wang, C. Ballesteros-Merino, S. M. Jensen, J. McGuire, M. Jiang, C. Coltharp, B. Remeniuk, I. Wistuba, D. Locke, E. R. Parra, B. A. Fox, D. L. Rimm and C. Hoyt, J Immunother Cancer, 2021, 9, e002197.

88 W. Huang, K. Hennrick and S. Drew, Human pathology, 2013, 44, 29–38.

89. P. W. Hamilton, P. Bankhead, Y. Wang, R. Hutchinson, D. Kieran, D. G. McArt, J. James and M. Salto-Tellez, Methods, 2014, 70, 59–73.

90. N. Goossens, S. Nakagawa, X. Sun and Y. Hoshida, Transl Cancer Res, 2015, 4, 256–269.

91. J. R. Lin, B. Izar, S. Wang, C. Yapp, S. Mei, P. M. Shah, S. Santagata and P. K. Sorger, Elife, 2018, 7, e31657.

92 A. J. Radtke, E. Kandov, B. Lowekamp, E. Speranza, C. J. Chu, A. Gola, N. Thakur, R. Shih, L. Yao, Z. R. Yaniv, R. T. Beuschel, J. Kabat, J. Croteau, J. Davis, J. M. Hernandez and R. N. Germain, Proc Natl Acad Sci U S A, 2020, 117, 33455–33465.

93. A. Somarakis, V. V. Unen, F. Koning, B. P. F. Lelieveldt and T. Höllt, IEEE Transactions on Visualization and Computer Graphics, 2019, 1–1.

94. N. Eling, N. Damond, T. Hoch and B. Bodenmiller, Bioinformatics, 2020, 36, 5706–5708.

95. D. Schapiro, A. Sokolov, C. Yapp, Y. A. Chen, J. L. Muhlich, J. Hess, A. L. Creason, A. J. Nirmal, G. J. Baker, M. K. Nariya, J. R. Lin, Z. Maliga, C. A. Jacobson, M. W. Hodgman, J. Ruokonen, S. L. Farhi, D. Abbondanza, E. T. McKinley, D. Persson, C. Betts, S. Sivagnanam, A. Regev, J. Goecks, R. J. Coffey, L. M. Coussens, S. Santagata and P. K. Sorger, Nat Methods, 2022, 19, 311–315.

96. T. M. Ashhurst, F. Marsh-Wakefield, G. H. Putri, A. G. Spiteri, D. Shinko, M. N. Read, A. L. Smith and N. J. C. King, Cytometry A, 2022, 101, 237–253.

97. W. C. C. Tan, S. N. Nerurkar, H. Y. Cai, H. H. M. Ng, D. Wu, Y. T. F. Wee, J. C. T. Lim, J. Yeong and T. K. H. Lim, Cancer Commun (Lond), 2020, 40, 135–153.

98. C. P. Hans, D. D. Weisenburger, T. C. Greiner, R. D. Gascoyne, J. Delabie, G. Ott, H. K. Muller-Hermelink, E. Campo, R. M. Braziel, E. S. Jaffe, Z. Pan, P. Farinha, L. M. Smith, B. Falini, A. H. Banham, A. Rosenwald, L. M. Staudt, J. M. Connors, J. O. Armitage and W. C. Chan, Blood, 2004, 103, 275–282.

99. W. W. Choi, D. D. Weisenburger, T. C. Greiner, M. A. Piris, A. H. Banham, J. Delabie, R. M. Braziel, H. Geng, J. Iqbal, G. Lenz, J. M. Vose, C. P. Hans, K. Fu, L. M. Smith, M. Li, Z. Liu, R. D. Gascoyne, A. Rosenwald, G. Ott, L. M. Rimsza, E. Campo, E. S. Jaffe, D. L. Jaye, L. M. Staudt and W. C. Chan, Clin Cancer Res, 2009, 15, 5494–5502.

100. T. Aoki, A. Jiang, A. Xu, Y. Yin, A. Gamboa, K. Milne, K. Takata, T. Miyata-Takata, S. Wu, M. Warren, C. Strong, T. Goodyear, K. Morris, L. C. Chong, M. Hav, A. R. Colombo, A. Telenius, B. Merrill, S. Ben-Neriah, M. Power, A. S. Gerrie, A. P. Weng, A. Karsan, A. Roth, P. Farinha, D. W. Scott, K. J. Savage, B. H. Nelson, A. Merchant and C. Steidl, bioRxiv, 2023, DOI: 10.1101/2023.05.19.541331, 2023.2005.2019.541331.

101. M. P. Kumar, J. Du, G. Lagoudas, Y. Jiao, A. Sawyer, D. C. Drummond, D. A. Lauffenburger and A. Raue, Cell Rep, 2018, 25, 1458–1468 e1454.

102. R. Browaeys, W. Saelens and Y. Saeys, Nat Methods, 2020, 17, 159–162.

103. M. Efremova, M. Vento-Tormo, S. A. Teichmann and R. Vento-Tormo, Nat Protoc, 2020, 15, 1484–1506.

104. F. Noel, L. Massenet-Regad, I. Carmi-Levy, A. Cappuccio, M. Grandclaudon, C. Trichot, Y. Kieffer, F. Mechta-Grigoriou and V. Soumelis, Nat Commun, 2021, 12, 1089.

105. N. F. Greenwald, G. Miller, E. Moen, A. Kong, A. Kagel, T. Dougherty, C. C. Fullaway, B. J. McIntosh, K. X. Leow, M. S. Schwartz, C. Pavelchek, S. Cui, I. Camplisson, O. Bar-Tal, J. Singh, M. Fong, G. Chaudhry, Z. Abraham, J. Moseley, S. Warshawsky, E. Soon, S. Greenbaum, T. Risom, T. Hollmann, S. C. Bendall, L. Keren, W. Graf, M. Angelo and D. Van Valen, Nat Biotechnol, 2022, 40, 555–565.

106. C. Sommer, C. Straehle, U. Köthe and F. A. Hamprecht, 2011.

107. M. Sorin, M. Rezanejad, E. Karimi, B. Fiset, L. Desharnais, L. J. M. Perus, S. Milette, M. W. Yu, S. M. Maritan, S. Doré, É. Pichette, W. Enlow, A. Gagné, Y. Wei, M. Orain, V. S. K. Manem, R. Rayes, P. M. Siegel, S. Camilleri-Broët, P. O. Fiset, P. Desmeules, J. D. Spicer, D. F. Quail, P. Joubert and L. A. Walsh, Nature, 2023, 614, 548–554.

108. E. Karimi, M. W. Yu, S. M. Maritan, L. J. M. Perus, M. Rezanejad, M. Sorin, M. Dankner, P. Fallah, S. Doré, D. Zuo, B. Fiset, D. J. Kloosterman, L. Ramsay, Y. Wei, S. Lam, R. Alsajjan, I. R. Watson, G. Roldan Urgoiti, M. Park, D. Brandsma, D. L. Senger, J. A. Chan, L. Akkari, K. Petrecca, M.-C. Guiot, P. M. Siegel, D. F. Quail and L. A. Walsh, Nature, 2023, 614, 555–563.

109. E. Lubeck and L. Cai, Nature Methods, 2012, 9, 743–748.

110. P. L. Ståhl, F. Salmén, S. Vickovic, A. Lundmark, J. F. Navarro, J. Magnusson, S. Giacomello, M. Asp, J. O. Westholm, M. Huss, A. Mollbrink, S. Linnarsson, S. Codeluppi, Å. Borg, F. Pontén, P. I. Costea, P. Sahlén, J. Mulder, O. Bergmann, J. Lundeberg and J. Frisén, Science, 2016, 353, 78–82.

111. C. R. Merritt, G. T. Ong, S. E. Church, K. Barker, P. Danaher, G. Geiss, M. Hoang, J. Jung, Y. Liang, J. McKay-Fleisch, K. Nguyen, Z. Norgaard, K. Sorg, I. Sprague, C. Warren, S. Warren, P. J. Webster, Z. Zhou, D. R. Zollinger, D. L. Dunaway, G. B. Mills and J. M. Beechem, Nature Biotechnology, 2020, 38, 586–599.

112. E. Gracia Villacampa, L. Larsson, R. Mirzazadeh, L. Kvastad, A. Andersson, A. Mollbrink, G. Kokaraki, V. Monteil, N. Schultz, K. S. Appelberg, N. Montserrat, H. Zhang, J. M. Penninger, W. Miesbach, A. Mirazimi, J. Carlson and J. Lundeberg, Cell Genomics, 2021, 1, 100065.

113. A. Chen, S. Liao, M. Cheng, K. Ma, L. Wu, Y. Lai, X. Qiu, J. Yang, J. Xu, S. Hao, X. Wang, H. Lu, X. Chen, X. Liu, X. Huang, Z. Li, Y. Hong, Y. Jiang, J. Peng, S. Liu, M. Shen, C. Liu, Q. Li, Y. Yuan, X. Wei, H. Zheng, W. Feng, Z. Wang, Y. Liu, Z. Wang, Y. Yang, H. Xiang, L. Han, B. Qin, P. Guo, G. Lai, P. Muñoz-Cánoves, P. H. Maxwell, J. P. Thiery, Q.-F. Wu, F. Zhao, B. Chen, M. Li, X. Dai, S. Wang, H. Kuang, J. Hui, L. Wang, J.-F. Fei, O. Wang, X. Wei, H. Lu, B. Wang, S. Liu, Y. Gu, M. Ni, W. Zhang, F. Mu, Y. Yin, H. Yang, M. Lisby, R. J. Cornall, J. Mulder, M. Uhlén, M. A. Esteban, Y. Li, L. Liu, X. Xu and J. Wang, Cell, 2022, 185, 1777–1792.e1721.

114. F. C. Chan, A. Mottok, A. S. Gerrie, M. Power, M. Nijland, A. Diepstra, A. van den Berg, P. Kamper, F. d’Amore, A. L. d’Amore, S. Hamilton-Dutoit, K. J. Savage, S. P. Shah, J. M. Connors, R. D. Gascoyne, D. W. Scott and C. Steidl, J Clin Oncol, 2017, 35, 3722–3733.

115. J. H. Friedman, T. Hastie and R. Tibshirani, Journal of Statistical Software, 2010, 33, 1–22.

116. N. Simon, J. H. Friedman, T. Hastie and R. Tibshirani, Journal of Statistical Software, 2011, 39, 1–13.

117. W. N. Venables and B. D. Ripley, Modern applied statistics with S-PLUS, Springer Science & Business Media, 2013.

