## Supplemental Figures for "Single Cell Spatial Analysis and Biomarker Discovery in Hodgkin Lymphoma"

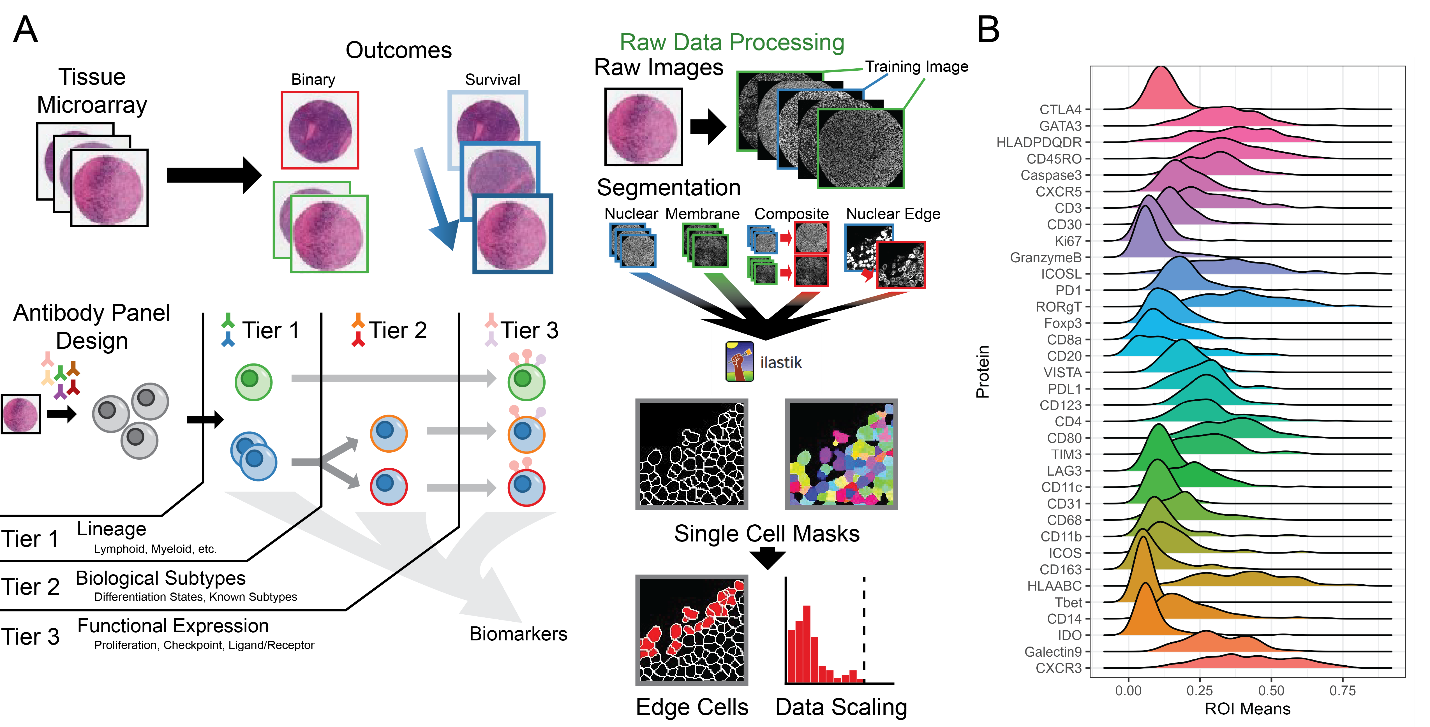
Supplementary Figure 1 Caption, associated with Figure 1. IMC Pipeline and Summary. A. For each ROI obtained from the TMA, clinical data such as patient outcomes and single cell data were recorded. Outcomes were stored as binary or categorical classes (EBV status, +/- 1 year relapse), or survival. The antibody panel was designed for hierarchical clustering using lineage markers into major cell phenotypes (Tier 1), followed by additional clustering into cell subtypes (Tier 2) and functional subtypes (Tier 3). Cell classifications generated using the first or second clustering steps were developed into biomarker candidates. Single cell data were generated by segmentation using the ilastik pipeline, with edge cells during spatial analysis as having biased spatial patterns. B. Following data scaling, mean expression of each marker per ROI was approximately normally distributed.

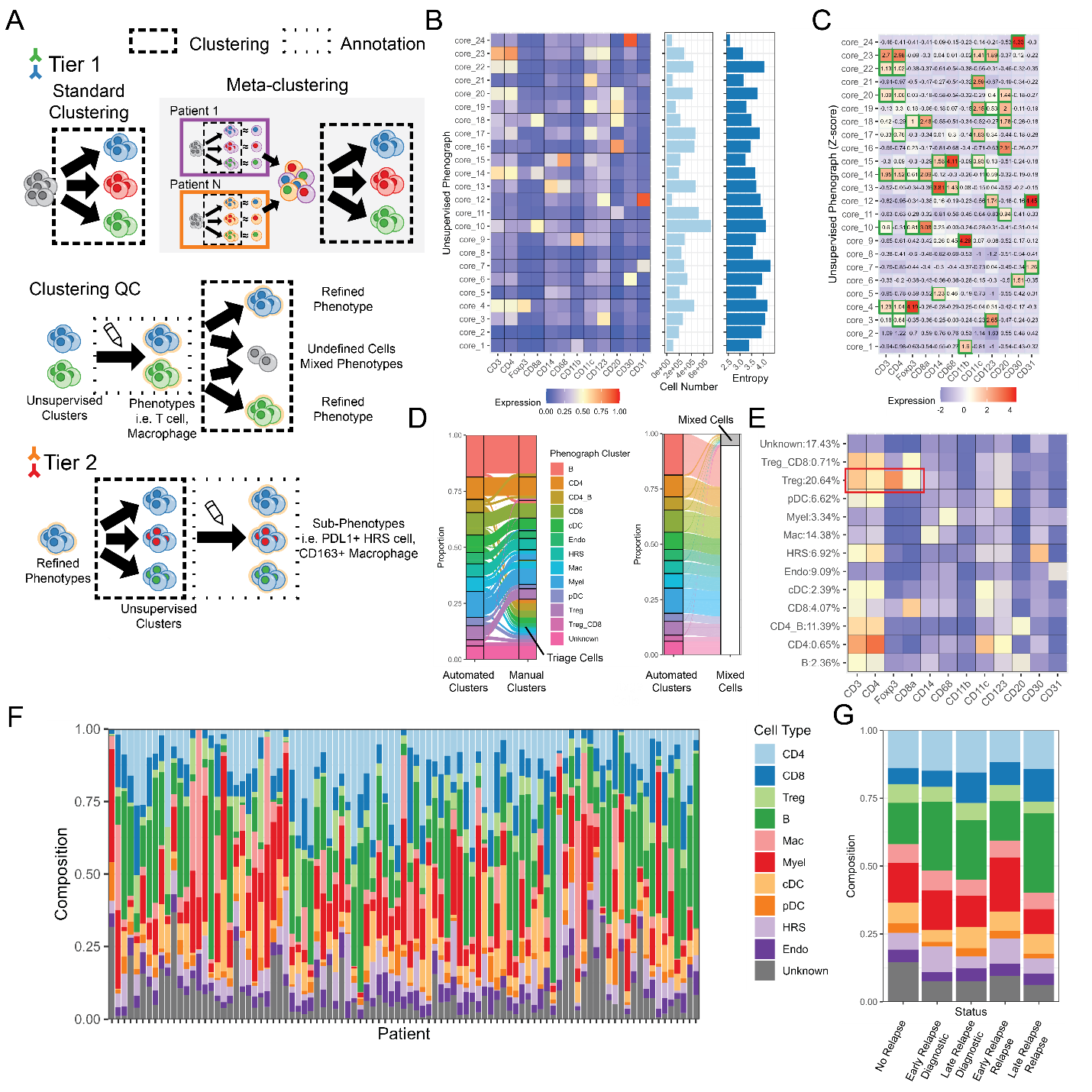
Supplemental Figure 2 Caption, associated with Figure 2. Phenotyping pipeline. A. A hierarchical clustering pipeline was used to perform cell phenotyping and extract cell types and subtypes of interest. Both automated clustering steps (dashed boxes) and manual annotations (dotted boxes) were used. Depending on the complexity of the data, a two-step metaclustering step can be used initially. The results of the pipeline are major cell type and cell subtype labels assigned to each cell, with multiple labels allowed per cell and ambiguous cells labeled “Unknown”. B. A heatmap of the first clusters generated and their proportions. Entropy describes the diversity of patients containing each of the clusters, with low cluster entropy indicating that fewer patient samples contained cells of that cluster. C. Z-score transformation of mean cluster expression was used to automatically label cell types. D. An alluvial plot (left) shows the number of cells of each automatically labeled cell type that was able to be manually assigned to a major cell type. Triage cells were those expressing ambiguous combinations of proteins. A second alluvial plot shows the number of cells in each automated cluster that were ultimately classified as “mixed”, likely representing overlapping cells in space. E. Mixed cells present combinations of well-defined subtypes, as shown in a heatmap (CD4+/CD8+/Treg mixed cell in red). F. Per patient distributions of each cell type. G. Cell type distributions divided by patient relapse status.

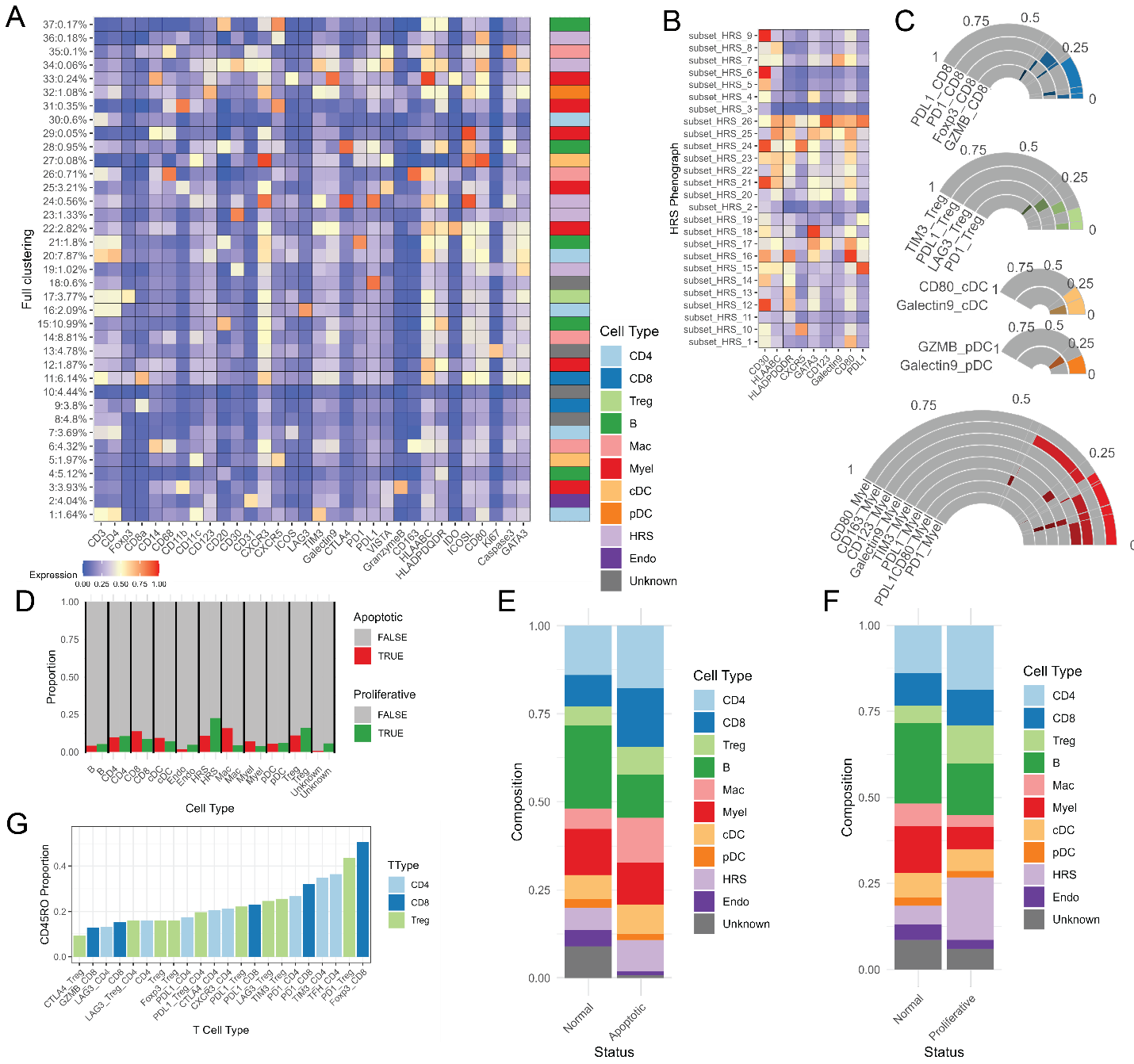
Supplemental Figure 3 Caption, associated with Figure 2. Cell subtype phenotyping. A. A single clustering step of all relevant proteins generated similar clusters as the metaclustering approach. B. HRS-specific clustering using HRS-specific markers. C. Relative proportions of CD8+, Treg, cDC, pDC, and myeloid subtypes expressing relevant proteins, depending on the proteins available in the panel. D. The percentage of Ki67+ and Caspase3+ cells of each major cell type. E,F. Among proliferative (Ki67+) or apoptotic (Caspase3+) cells, the relative distribution of cell types. G. The percentage of CD45RO+ cells of each T cell subtype.

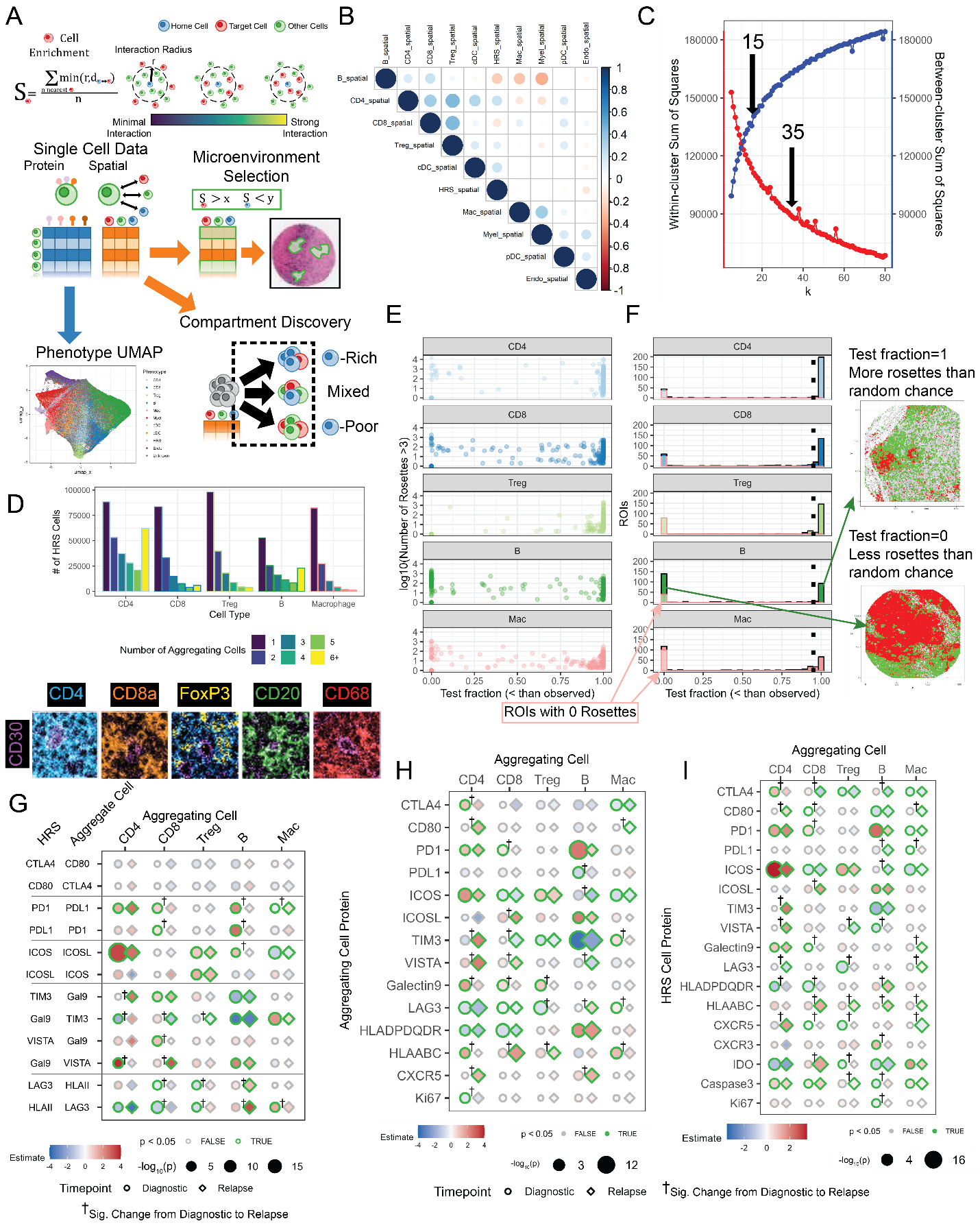
Supplemental Figure 4 Caption, associated with Figure 3. Spatial Protein Expression. A. A spatial enrichment metric was calculated using the 5-nearest neighbor average distance, modified by capping at 50 μm to remove distant cell effects. Nearest neighbor distance was similarly capped at 50 μm. Spatial data was treated similarly to protein data to generate clusters of cells with similar spatial environments by clustering, or to select cells based on thresholded spatial measurements. B. A correlation plot shows the relative enrichments between all cell types. C. The within-cluster and between-cluster distances for k-means clustering, with no clear elbow. D. The number of HRS cells found with each aggregate size for CD4+, CD8+, Treg, B, and macrophage cells, and sample images of large aggregates of each type. E. A plot of the number of aggregates in an ROI vs the test fraction of random replacement tests with more aggregates than expected. F. The histogram of test fractions among ROIs, with ROIs with no aggregates observed highlighted. G. Receptor-ligand coexpression measurements associated with aggregate size in relapse samples, with differences from diagnostic samples indicated by †. H. Mean expression of proteins on aggregating cells associated with aggregate size in diagnostic and relapse samples. I. Expression of proteins on HRS cells associated with aggregate size in diagnostic and relapse samples.

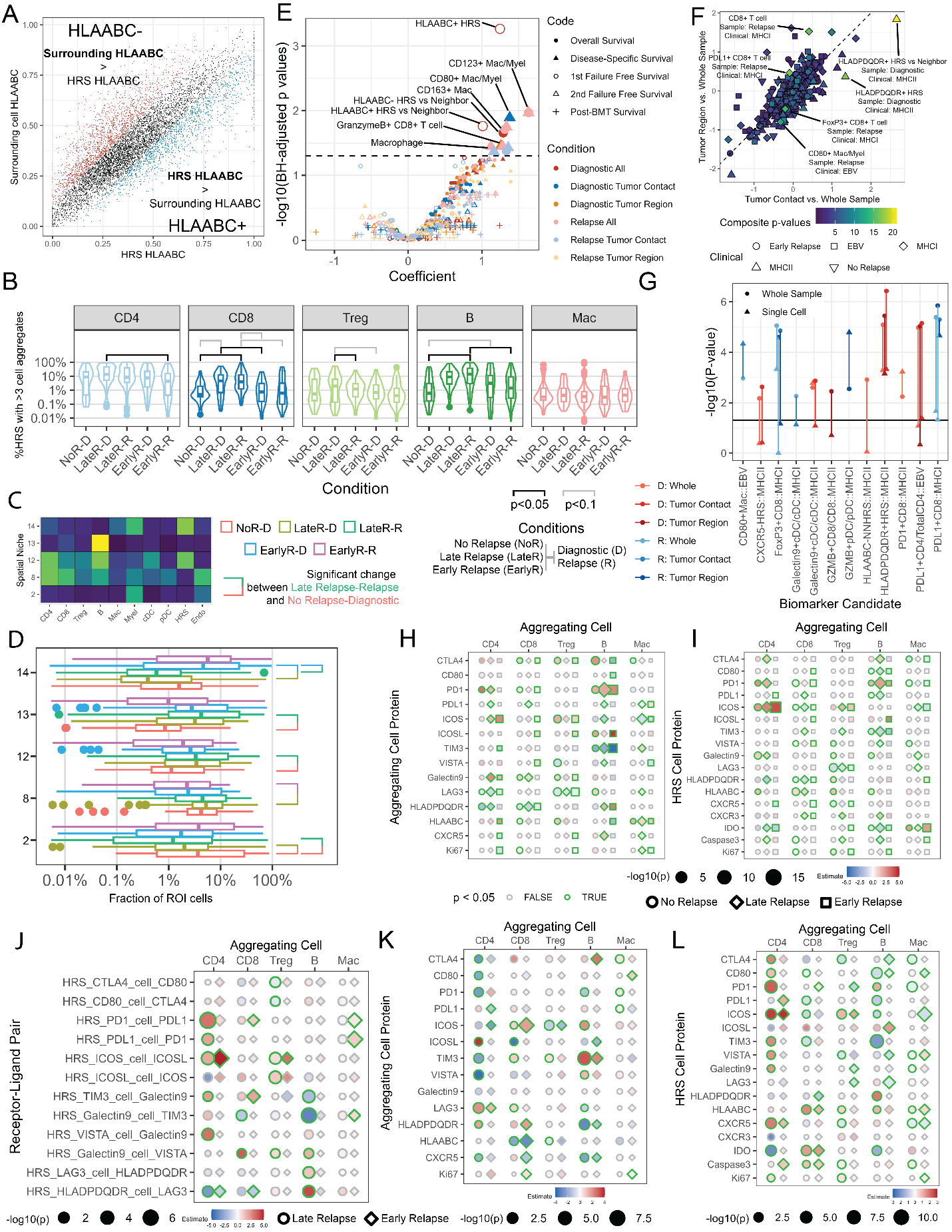
Supplementary Figure 5 Caption, associated with Figure 4. Biomarker Testing. A. HRS HLAABC and surrounding cell HLAABC used to reproduce the MHC- HRS cell biomarker. B. Biomarker candidates significance by cell subsets for 5 types of survival analyses (OS - overall survival, PFS - progression free survival, FFS1 - first failure free survival, FFS2 - second failure free survival, BMTFFS - post bone marrow transplant first failure free survival). Multiple test correction by Benjamini-Hochberg FDR. C. Biomarker candidate relative significance for categorical variables. D. Aggregation frequency vs. relapse status and timepoint, significance measured by Tukey’s HSD. E. The spatial composition of 5 niches with significant abundance differences between clinical statuses are shown. F. The abundances and statistical differences (Tukey’s HSD) of the 5 spatial niches are shown. G. Single cell biomarker candidate comparisons, D = Diagnostic, R = Relapse. H-I. Relapse status-dependent protein expression associated with aggregate size on aggregating cells or HRS cells in diagnostic samples. J-L. Differences observed in receptor-ligand coexpression, aggregating cell protein, and HRS cell protein expression between early and late relapses in relapse samples.

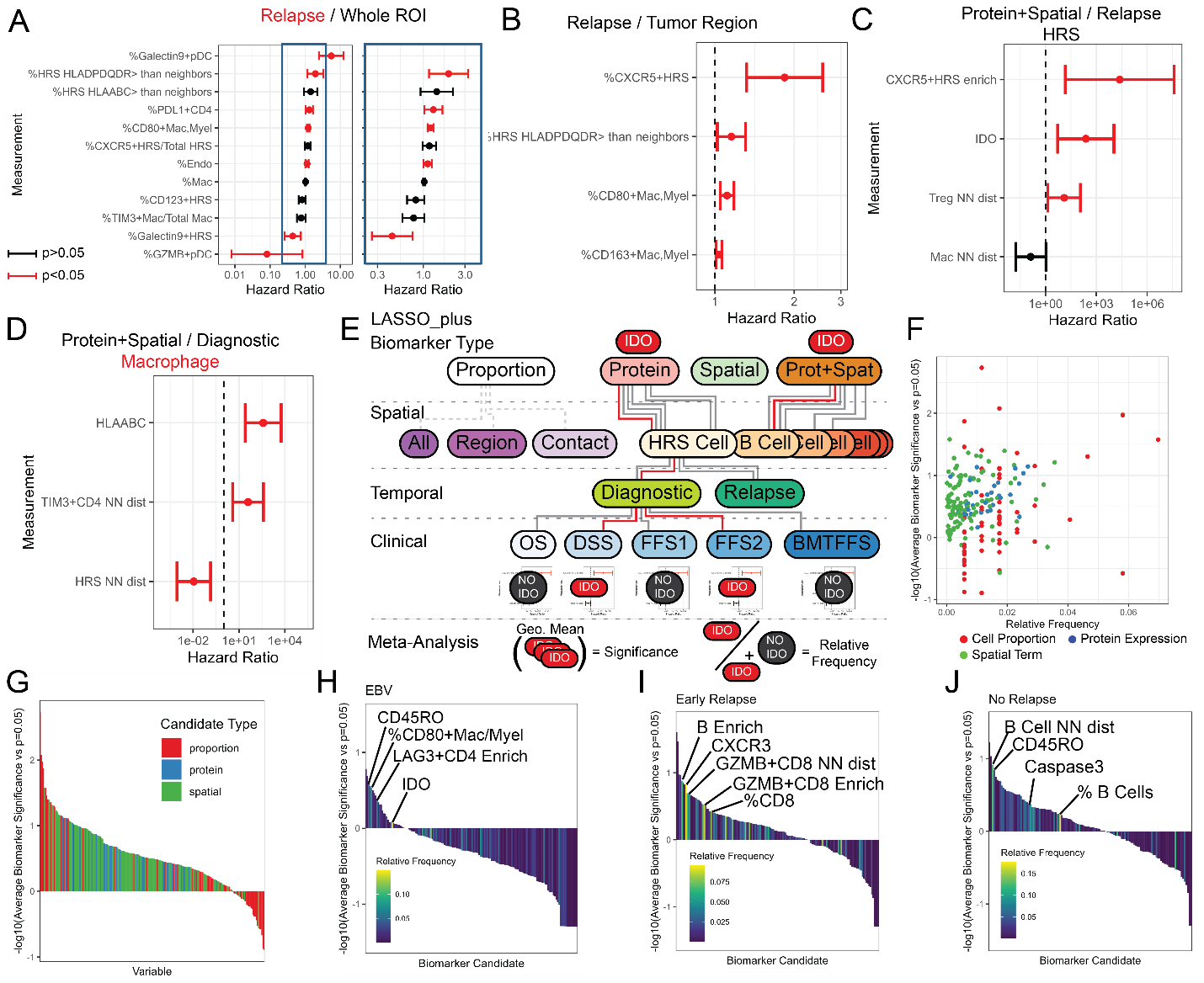
Supplemental Figure 6 Caption, associated with Figure 5. Biomarker Discovery. A. LASSO_plus output of biomarker candidate set of cell proportions for relapse samples with zoom in inset. B. Biomarker candidate set for relapse samples in tumor contact digital biopsy. C. Biomarker candidate set of protein and spatial measurements of HRS cells. D. Biomarker candidate set for proteins and spatial measurements on macrophages in diagnostic samples. E. Biomarker meta-analysis was performed using LASSO_plus on combinations of biomarker types, spatial, temporal, and clinical conditions. For each biomarker candidate, the number of times it was selected by LASSO_plus was compared to the maximum possible times it could be selected to obtain the biomarker relative frequency. The geometric mean of p-values was also calculated. F. The relative frequency and geometric mean of each biomarker candidate is plotted. G. Waterfall plot of biomarker candidates ranked by their relative significance vs p=0.05 and colored by the biomarker candidate type. H-J. Waterfall plot for biomarker candidates EBV status, early relapse, and no relapse.
